# ZNF865 (BLST) a Novel Regulator of DNA Damage and Cell Senescence in Back Pain

**DOI:** 10.1101/2025.06.13.659603

**Authors:** Christian Lewis, Hunter Levis, Jonah Holbrook, Jacob T. Polaski, Timothy D. Jacobsen, Josh Mizels, Molly Czachor, Sarah E. Gullbrand, Brian Diekman, James C. Iatridis, Jason Gertz, Brandon Lawrence, Robby D. Bowles

## Abstract

Senescence has been shown to contribute to the progression of aging related diseases including degenerative disc disease (DDD). However, the mechanisms regulating senescence in the intervertebral disc (IVD) and other tissues/diseases remain poorly understood. Recently, ZNF865 (BLST) was identified as a previously uncharacterized zinc finger protein, that regulates a wide array of genes related to protein processing, cell senescence and DNA damage repair. Here, we show that ZNF865 expression decreases with age and pathology in human and mouse IVD samples. Utilizing CRISPR-guided gene modulation, we show that ZNF865 is necessary for healthy cell function and is a critical protein in regulating senescence and DNA damage in IVD cells, with implications for a wide range of tissues and organs. We also demonstrate that downregulation of ZNF865 induces senescence in asymptomatic human nucleus pulposus (NP) cells. Conversely, restoration in degenerative NP cells mitigates senescence, SASP expression and DNA damage, enhances ECM anabolism, and restores chromatin accessibility and gene expression to a healthy state. In vivo, CRISPRi of ZNF865 induces disc degeneration and painful behaviors. Collectively, our findings establish ZNF865 as a novel regulator of genome stability and a potential therapeutic target for mediating senescence/DNA damage in aging related diseases.

## Introduction

Cellular senescence has been increasingly recognized as a fundamental driver of disease progression in aging related diseases. During senescence, cells undergo stable cell cycle arrest often described as a protective mechanism against oncogenic transformation and excessive DNA damage^1,2^. These cells secrete many pro-inflammatory catabolic factors collectively known as the senescence associated secretory phenotype (SASP) that can disrupt the tissue microenvironment and further sustain the inflammatory environment through complex positive feedback loops^2–4^. In many aging related disorders, the increased presence of senescent cells is thought to be a major contributor to disease progression. In the central nervous system, the accumulation of senescent glial cells has been shown to contribute to inflammation and neurodegeneration potentially accelerating diseases such as Alzheimer’s^5,6^ and Parkinson’s^7^. In the lungs, senescent fibroblast populations are associated with impaired repair mechanisms and fibrotic remodeling in pulmonary fibrosis^8,9^. In the musculoskeletal system, senescent chondrocytes and other cell types of the joint have been shown to contribute to cartilage matrix breakdown and synovial inflammation in osteoarthritis^10^ as well as increased inflammation and matrix breakdown in degenerative intervertebral discs (IVD)^11^. *Copp et al.* showed that chondrocytes obtained from femur cartilage of osteoarthritic patients had significantly higher levels of DNA damage, but also that the amount of DNA damage present in healthy chondrocytes increased with age^12^, indicating that excessive DNA damage may be an indicator of cells capable of contributing to disease pathology in osteoarthritis, and other aging-related musculoskeletal disorders. Despite this increased attention on DNA damage and senescence in aging related disorders, the underlying molecular mechanisms are not completely understood and this manuscript introduces a novel gene/protein regulator of these processes, investigating it here specifically in back pain.

Back pain is one of the most common musculoskeletal disorders and a leading cause of disability worldwide^13^, with a large population of people expected to experience it at some point in their lifetime^14,15^. Chronic back pain is often debilitating, with 10% of individuals with back pain reportedly being forced to change jobs or stop working entirely due to severe pain^16^. Back pain incidence and severity have also been shown to increase with age with 40% of back pain attributed to degenerative disc disease (DDD)^17^. While back pain etiology is diverse and multifactorial, growing evidence suggests biological aging and senescence may be major contributing factors. Traditionally DDD onset is associated with the breakdown of the disc tissue due to genetics^18^, aging^19,20^, loading history^21^ and inadequate metabolite transport^22^. However, as DDD progresses, studies have shown there is decreased cell activity due to increased cell senescence^23,24^. Additionally, degenerative discs are often characterized by increased presence of pro-inflammatory cytokines, proteases and pain related growth factors consistent with the SASP phenotype. These changes to extracellular matrix composition can lead to mechanical instability, pain and disability rates^22,25–27^. As a result, senescence represents an emerging potential therapeutic target for back pain patients, but the literature in this space is limited, and more work understanding the molecular mechanism of senescence in back pain is needed.

Targeting senescent cells via senolytics within the IVD has provided promising results in reducing the release of SASP factors in isolated IVD cells and intact human IVDs^28,29^. Oral administration of o-vanillin and RG-7112 resulted in decreased pain and histological degeneration in the IVD of sparc-null mice^30^. Additionally, intra-peritoneal delivery of the senolytics dasatinib and quercetin from 14 to 23 months of age significantly improved nucleus pulposus (NP) and annulus fibrosus outcomes in wild type mice^31^. Overall, this work further distinguishes cellular senescence as an important mediator of IVD degeneration and highlights the critical need to develop novel tools that target senescence directly to prevent degeneration and reduce pain caused by SASP.

Recently, we identified ZNF865, also known as BLST, a gene with no known functional role, and no studies on its molecular actions, and utilized it as a cell engineering tool to enhance the tissue deposition and mechanical properties of engineered IVDs^32^. Initially we identified this gene in multiple different genome wide CRISPR perturbations screens^33,34^ for cell engineering applications. However, our RNA sequencing data also linked ZNF865 to DNA damage and senescence^32^. At the time, only bioinformatic information was available that identified ZNF865 as 1059 amino acids long, as a C2H2 zinc finger (ZNF) protein, and containing six disordered regions, twenty unique ZNF domains, two transactivation domains (TADs) and two TGEKP linkers (**Figure 1A**)^35^. The presence of the two TADs indicate that ZNF865 likely stimulates transcriptional activity by interacting with transcriptional regulators^36^. Despite broad expression across all cell types (**Figure 1B**)^35,37^, there are no publications functionally characterizing the biological role of ZNF865. Additionally, a rare disease mutation in ZNF865 is associated with a wide range of symptoms including global developmental delay, joint hypermobility, muscle hypertonia, loose inelastic skin, hand tremors, and eye/ear/brain abnormalities indicating effects across a wide range of tissues and organs in human patients^38^, indicating a clear important biological role for this gene/protein. To this end, with the growing link between senescence and back pain, and ZNF865s link to DNA damage and senescence, we investigated ZNF865 in the context of back pain within human back pain patient samples and multiple rodent models. This represents the first data on the functional role of ZNF865 in any disease and required us to start with no existing tools to study it.

**Figure 1:**
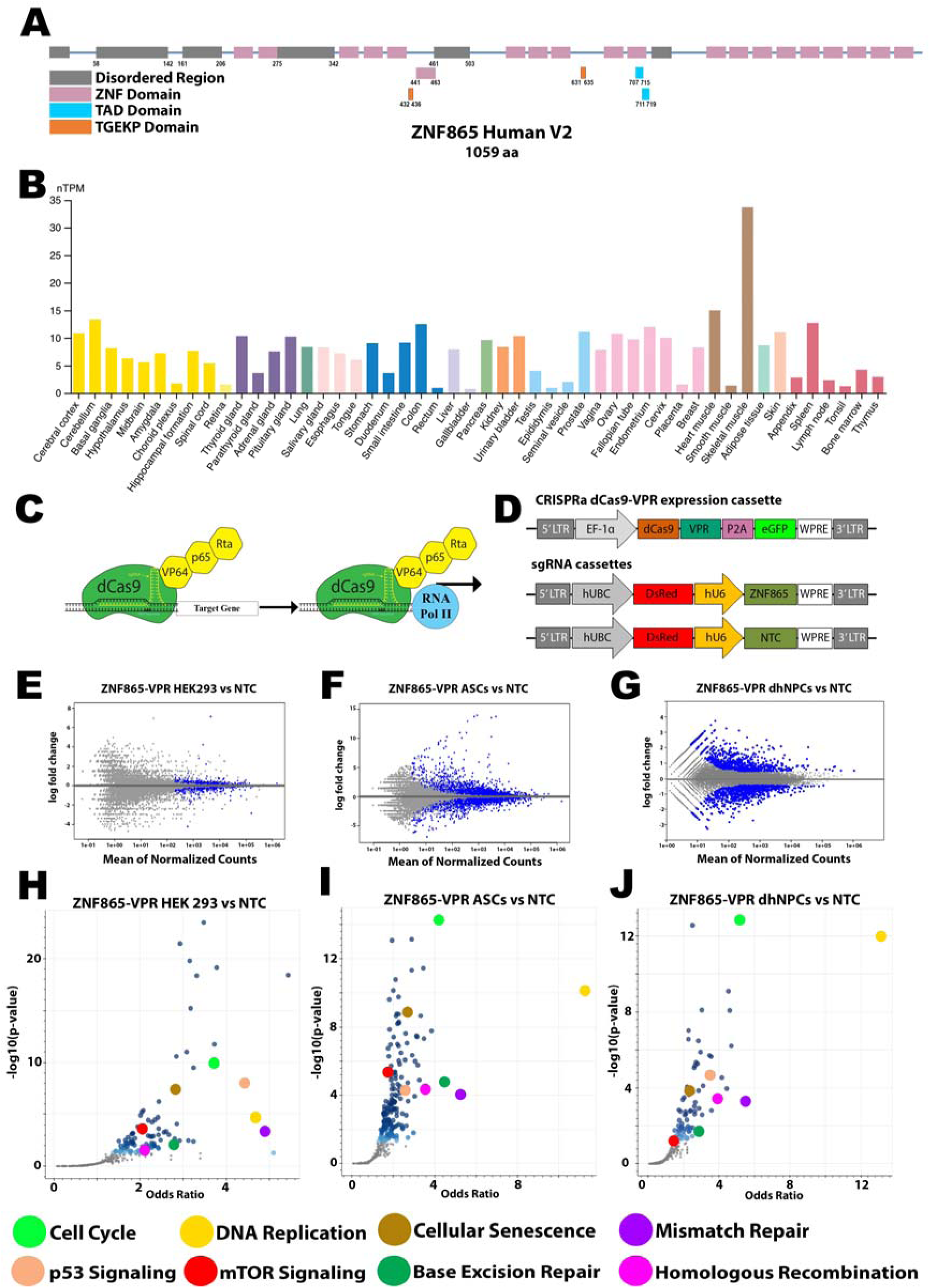
ZNF865 Regulates Similar Gene Networks Across Multiple Cell Types. **A** Structure of the human ZNF865 gene. **B** ZNF865 expression across different human tissue sources. nTPM = normalized transcripts per million. **C** dCas9-VPR CRISPRa system and **D** plasmid maps for lentiviral delivery targeting ZNF865 or NTC. **E** MA plots from RNA-seq on ZNF865 upregulated HEK293 cells, **F** ASCs and **G** human dhNPCs. **H** Volcano plots showing the top common molecular process affected by ZNF865 upregulation across HEK293 cells, **I** ASCs and **J** human dhNPCs using the KEGG 2021 Human Pathway. *NTC is non­target control.*

Here we link ZNF865 as an important driver of discogenic back pain through the regulation of senescence and DNA damage. This investigation further elucidated the role DNA damage plays in the onset and perpetuation of the degenerative phenotype in DDD indicating that the depletion of ZNF865 is sufficient to induce DNA damage, disc degeneration and painful phenotypes both *in vitro* and *in vivo*. Importantly, upregulation of ZNF865 is capable of reprogramming human disc cells from back pain patients to return them to a healthier phenotype. Prior to this work, ZNF865 remained largely uncharacterized, addressing a significant gap in literature by providing the first functional characterization of its role in disc cell homeostasis, senescence and DNA damage. Furthermore, these insights identify ZNF865 as a key modulator in maintaining genomic stability and as a potential therapeutic target in other aging related diseases hallmarked by senescence.

## Results

### ZNF865 upregulation alters related cell functions across multiple cell types

To begin investigating the molecular role of ZNF865, CRISPR activation (CRISPRa) was used to increase the expression of ZNF865 using the dCas9-VPR system alongside a non-target control (**Figure 1C,1D**). RNAseq was then performed on three different human cell types; human embryonic kidney cells (HEK293), adipose derived stem cells (ASCs) and degenerative human NP cells (dhNPCs), and comparisons were made across the CRISPRa ZNF865 and non-target samples (**Figure 1E, 1F, 1G**). Upregulation of ZNF865 in each of these cell types showed 2,699, 6,647 and 1,752 significantly differentially expressed genes, respectively. Utilizing Enrichr^39^, analysis of these genes showed a number of significantly differentially regulated molecular processes using the KEGG 2021 Human Pathway that were shared across cell types (**Figure 1H-1J, Table S2-S4**)^40^. The notable pathways across all three cell types included cell cycle, DNA replication, cellular senescence, mismatch repair, p53 signaling, mTOR signaling, base excision repair, and homologous recombination. The shared molecular processes suggested a role for ZNF865 in regulating cell division, DNA repair, and senescence.

### ZNF865 expression is correlated with human degenerative disc disease and aging

To evaluate whether ZNF865 expression is pathologically relevant within DDD, gene expression was measured via qRT-PCR on dhNPCs from patients undergoing surgical treatment for low back pain, and asymptomatic NP cells (hNPCs) from trauma patients at first passage after plating in vitro (**Figure 2A, Table S1**). We observed a strong correlation (R^2^=0.95, p=0.0054) between ZNF865 expression and age in hNPCs and significantly decreased expression in dhNPC samples at matched ages (**Figure 2B**). Immunofluorescence staining for ZNF865 matched the qRT-PCR data with significantly less ZNF865 staining in dhNPC samples compared to hNPCs (**Figure 2C,2D**). ZNF865 staining was localized to the nucleus of the cells, consistent with bioinformatic predictions that ZNF865 is a transcription factor. Of note, 28.7% of hNPCs stained for ZNF865, while only 16.8% of dhNPCs were stained, indicating that ZNF865 may not be expressed by all cells at a detectable level at a given time (**Figure 2E**).

**Figure 2:**
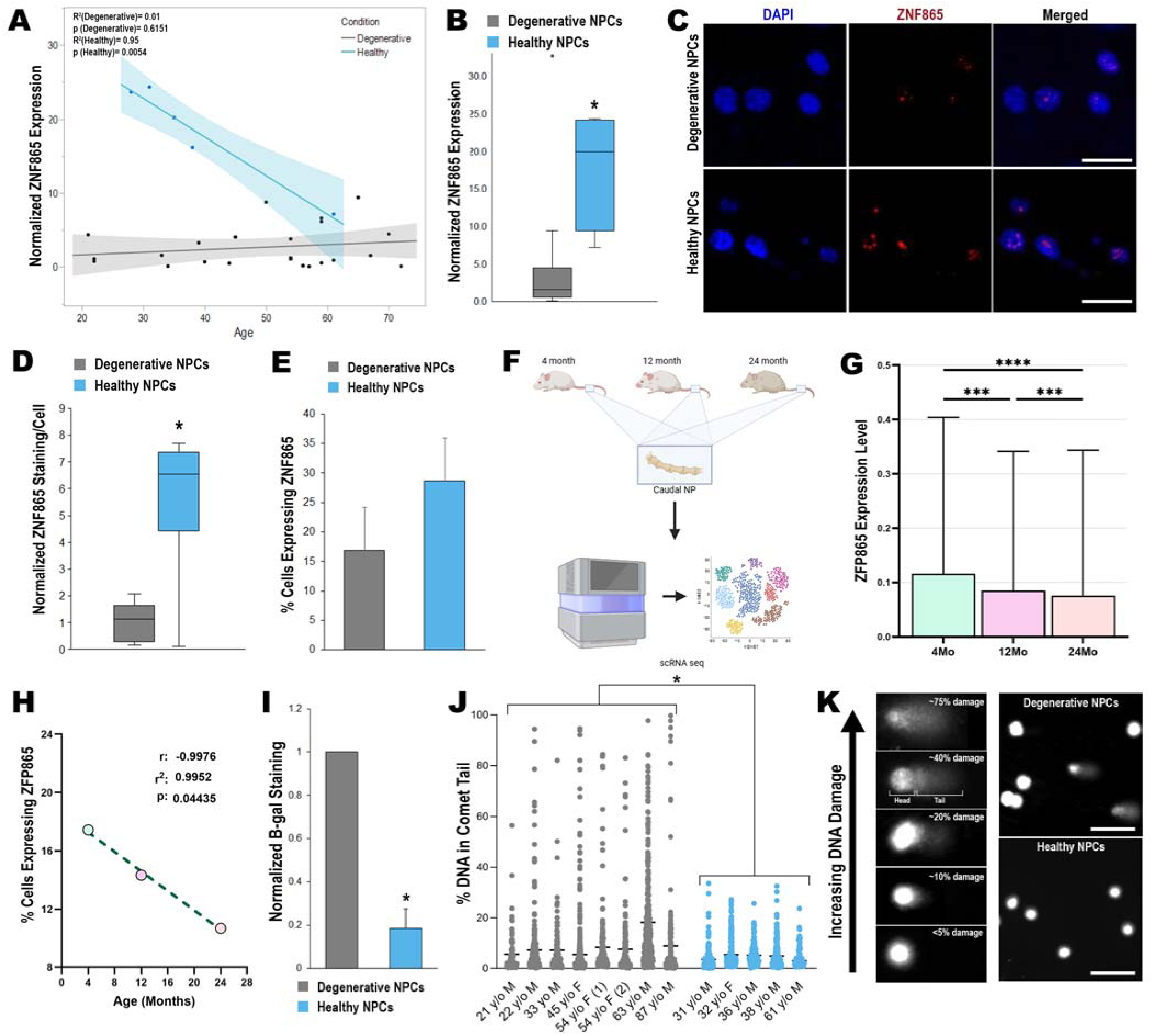
ZNF865 Expression is Correlated with Aging, Degeneration, and DNA Damage in Human and Mouse NP cells. **A** Normalized ZNF865 expression in hNPCs (n=5) and dhNPCs (n=23) in individual patients and **B** grouped by pathology. **C** Representative images from ZNF865 immunofluorescence staining (Blue=DAPI, Red=ZNF865). **D** ZNF865 immunofluorescence staining per cell normalized to the dhNPC level (n=7). **E** Percent of dhNPCs and hNPCs expressing ZNF865 from immunofluorescence staining (n=3-4). **F** Workflow for scRNA-seq in the caudal NP of aging mice. **G** ZFP865 expression level in the caudal NP of aging mice. **H** Percent of cells positive for ZFP865 in the aging mouse model. **I** Normalized P-galactosidase staining of healthy (n=4) and dhNPCs (n=5). **J** Percent DNA in the comet tail of naive hNPCs (n=5) and dhNPCs (n=8) measured by comet assay. **K** Images showing single cell comets with percent damage, and representative dhNPCs and hNPC examples. *(Scale bars = 50 pm. significance *=p<0.05, **=p<0.01, ***=p<0.001)*

To determine whether expression of this gene also changes in mouse IVD, ZFP865 expression (mouse orthologue of ZNF865)^41^ was examined in the caudal NP of mice at 4, 12 and 24 months of age (**Figure 2F**). These results were consistent with the human data, where ZFP865 expression decreased with increasing age. Both population average expression level (**Figure 2G**) and percentage of cells expressing (**Figure 2H**) showed decreases, with a strong negative correlation between cells expressing ZFP865 and age in the mouse (r = -0.9976, p = 0.04435).

Cell senescence has been proposed as a potential mechanism of DDD with degenerative discs showing increased levels of senescence (**Figure 2I)**^23,24,42^. However, the functional and molecular links between senescence and DDD are still relatively unknown. Given that our RNA-seq data indicates that both senescence and closely associated DNA repair pathways are significantly altered following CRISPRa ZNF865 upregulation (**Figure 1J**), we investigated DNA damage in the human dhNPCs and hNPCs using alkaline comet assays. Here we observed that the percentage of DNA damage in the comet tail in dhNPCs was significantly higher than in hNPCs (**Figure 2J, 2K**). Together, this data indicates a relationship between ZNF865 expression and DDD, as well as DDD progression and DNA damage. Moving forward we investigated the relationship between ZNF865 expression and cellular senescence and DNA damage in human NP cells.

### ZNF865 downregulation in asymptomatic human NP cells induces senescence and DNA damage

hNPCs were isolated from discarded surgical tissue from deidentified trauma patients (category 4 exemption as approved by the University of Utah IRB) and transduced with the CRISPRi vector targeting ZNF865 (ZNF865-KRAB) or a non-target control (**Figure 3A, Table S9**). Following transduction, cells were monitored for cell proliferation over 6 days. Samples transduced with ZNF865-KRAB showed decreased proliferative capacity compared to cells transduced with the non-target vector (**Figure 3B**). To determine if the reduction in cell growth observed in the ZNF865-KRAB samples was consistent with entry into senescence, hNPCs were stained for key markers of cellular senescence, including senescence associated β- galactosidase (β-gal), p16^INK4A^ (p16) and p21^waf^^1/Cip^^1^ (p21). The ZNF865-KRAB NP cells showed significantly higher levels of all three senescence markers, with 5.4-fold, 4.7-fold, and 6.9-fold increases in staining, respectively (**Figure 3C-3E, 4H**). Entry into senescence has commonly been linked to increased DNA damage as a way to prevent damaged cells from perpetuating their genomes in division^2,43^, and ZNF865 regulation had been demonstrated to regulate DNA damage repair pathways (**Figure 1**). To quantify the amount of DNA damage across patient samples, we performed alkaline comet assays 3 weeks after transduction. Here, the ZNF865-KRAB NP cells showed significantly higher levels of DNA damage compared to the non-target samples, indicating a functional connection between decreased ZNF865 expression and DNA damage (**Figure 3F,3G**). This data is consistent with the relationships observed between ZNF865, senescence and DNA damage in human patient NP cells (**Figure 2**) and our RNAseq data (**Figure 1**).

**Figure 3:**
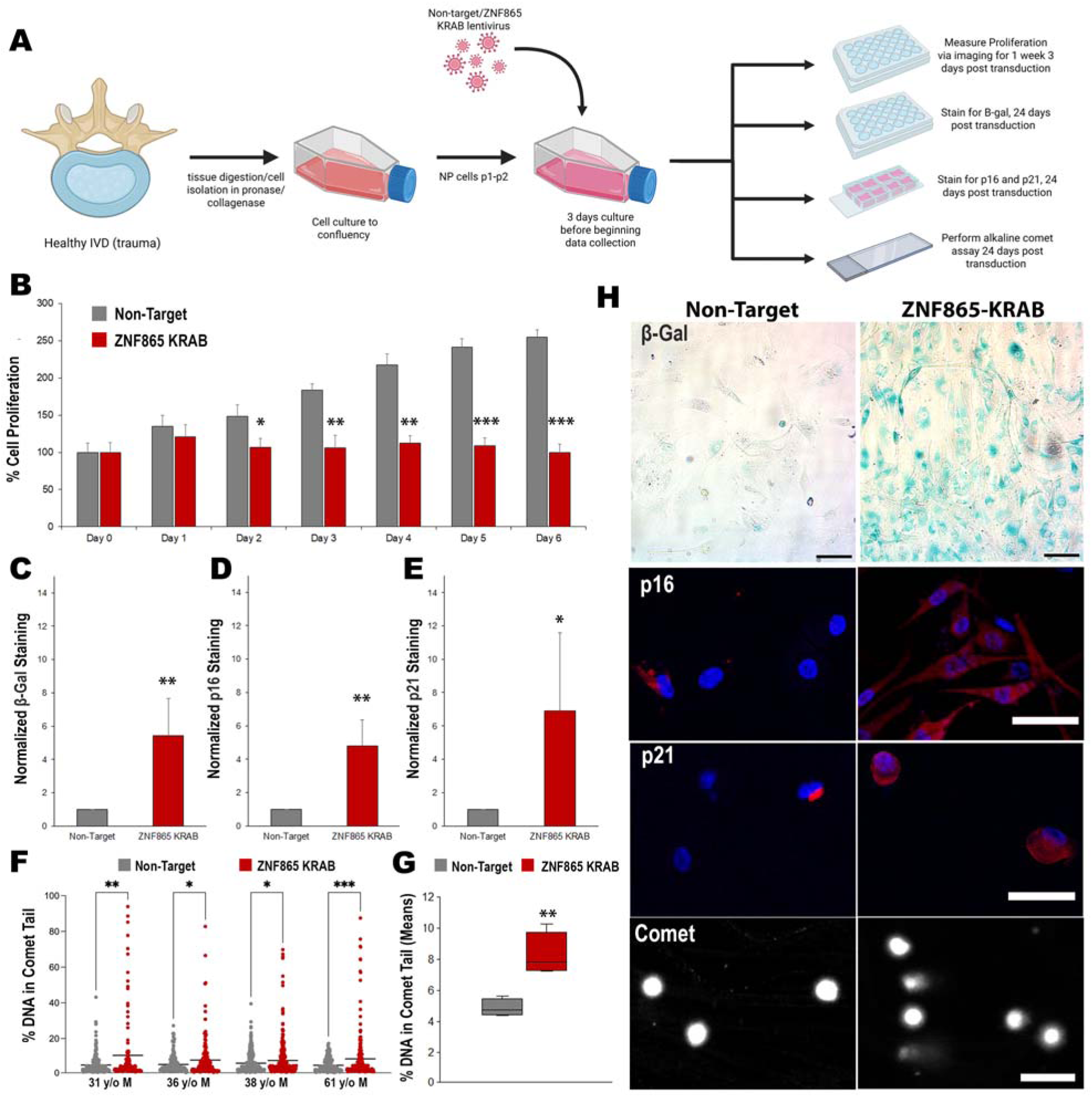
CRISPRi of ZNF865 decreases proliferation rates, increases senescence and increases DNA damage in hNPCs. **A** Workflow for CRISPRi of ZNF865 or non-target in hNPCs. **B** Quantified cell proliferation of healthy human non-target and ZNF865-KRAB NP cells (n=4). **C** SA-β-Galactosidase, **D** p16 and **E** p21 staining normalized to non-target (n=4). **F** Percent DNA in comet tail in individual patients obtained by comet assay. **G** Donor mean percent DNA in comet tail (n=4). **H** Representative images from SA-β-Galactosidase, p16 and p21 staining, and nuclei from comet assays. (Scale bar β-Gal = 100 *µm, scale bar others = 50 µm. Significance *=p<0.05, **=p<0.01, ***=p<0.0QP)*

### ZNF865 upregulation in degenerative human NP cells decreases senescence, SASP phenotype and DNA damage

To begin studying the effects of ZNF865 regulation in pathologic cells, dhNPCs were isolated from discarded surgical tissue from deidentified patients undergoing surgical treatment for chronic back pain. Cells were then transduced with ZNF865 upregulation cassettes (ZNF865-VPR) or a non-target control (**Figure 4A, Figure S1**). Following transduction, cells were monitored for cell proliferation via fluorescent imaging over 5 days. We observed significant increases in cell growth within 3 days in the ZNF865-VPR cells compared to the non-target controls (**Figure 4B**). This led to a significant decrease in doubling time in the ZNF865-VPR dhNPCs compared to the non-target dhNPC controls, at 2.49 days and 4.09 days respectively, with non-significant differences compared in the naïve hNPCs with a doubling time of 3.2 days (**Figure 4C**). We stained each treatment group for key senescence markers 10 days after transduction. Here, the non-target dhNPCs showed the highest levels of staining in β-gal, p16 and p21, with significantly lower levels of senescence staining in the ZNF865-VPR dhNPCs and hNPCs. Additionally, the ZNF865-VPR dhNPCs and hNPCs were non-significantly different from each other (**Figure 4D-4G**).

**Figure 4:**
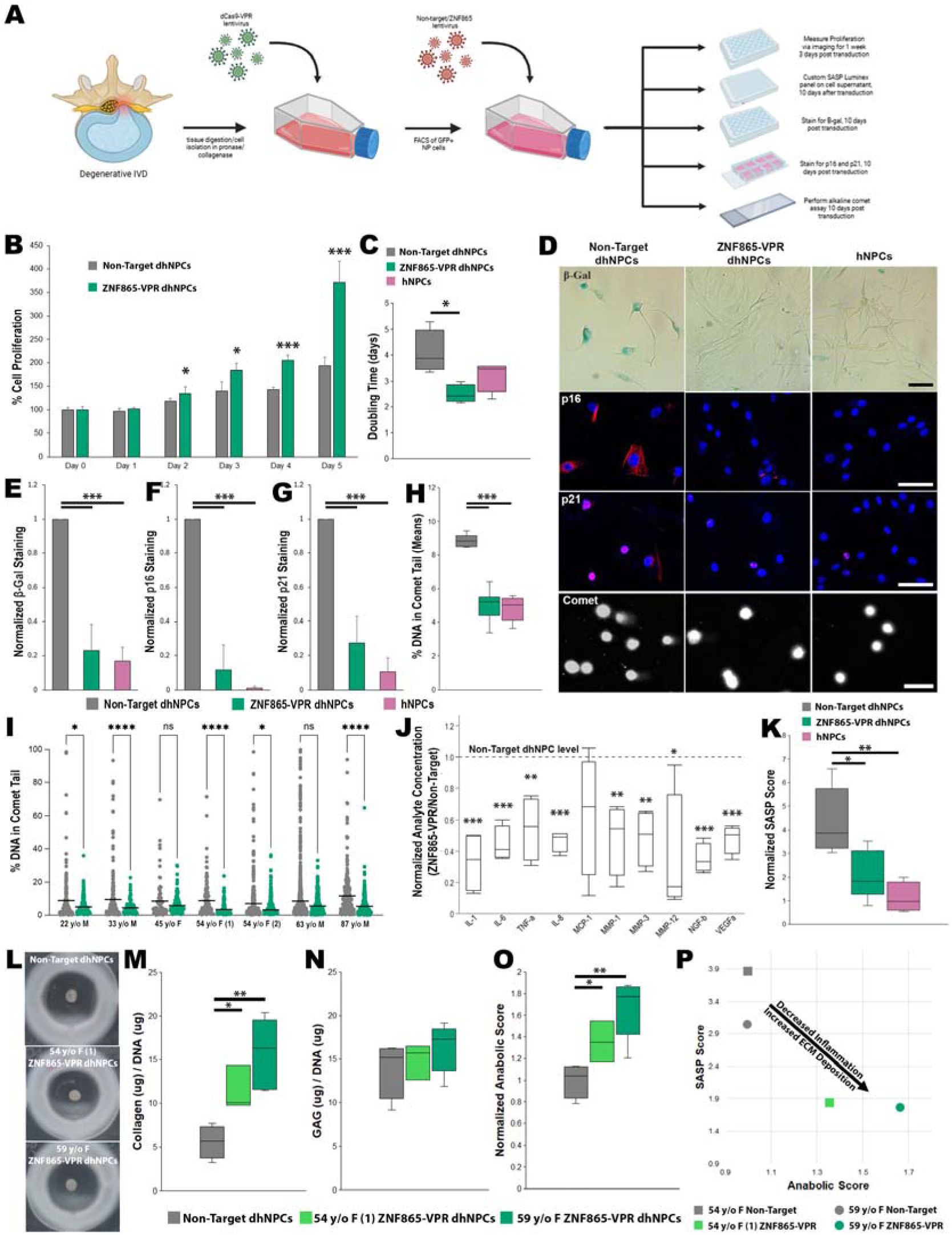
CRISPRa of ZNF865 restores proliferation, decreases senescence, decreases SASP phenotype and decreases DNA damage to healthy levels in dhNPCs. **A** Workflow for CRISPRa of ZNF865 or non-target in dhNPCs. **B** Quantified cell proliferation of human non-target dhNPCs and ZNF865-VPR dhNPCs (n=5). **C** Doubling times of non-target dhNPCs, ZNF865-VPR dhNPCs and hNPCs (n=5). **D** Representative images from P-Gal, p16 and p21 staining, and nuclei from comet assays. **E** SA-P-Galactosidase, **F** p16 and **G** p21 staining normalized to non-target (n=4-5). **H** Percent DNA in comet tail in non-target dhNPCs, ZNF865-VPR dhNPCs and hNPCs. **I** Donor mean percent DNA in comet tail (n=7). **J** Analyte concentration for SASP related inflammatory cytokines, proteases and proteins present in ZNF865-VPR dhNPCs normalized to non-target samples (n=4). Non-target levels indicated by dashed line. **K** SASP score normalized to hNPC values (n=4-5). **L** Representative images of pellet culture. **M** Total collagen production per DNA and **N** GAG production per DNA and **O** normalized anabolic scoring from pellet culture in pooled non-target dhNPCs and two ZNF865- VPR dhNPC patient samples (n=3-6). **P** Plot of showing relationship of SASP score and anabolic score after ZNF865 upregulation. Scale bars *= 50pm. Significance *=p<0.05, **=p<0.01, ***=p<0.00)*

To assess the ability to directly regulate DNA damage, we performed alkaline comet assays. Here we found that upregulation of ZNF865 significantly decreased the percentage of DNA in the comet tails in all patients, with five of the seven patients reaching significance (**Figure 4I**). When the mean damage of each donor is considered, there is a significant decrease in the ZNF865-VPR dhNPCs compared to the non-target controls, with the ZNF865-VPR dhNPCs matching the asymptomatic controls (**Figure 4H,4I**). This demonstrates ZNF865’s ability to directly regulate DNA damage and senescence observed in human samples from painful degenerative discs.

To better understand the downstream effects of reversing senescence and DNA damage in these cell populations, a Luminex assay was run on 10 SASP related proteins using the cell culture supernatant from the ZNF865-VPR dhNPCs, non-target control dhNPCs, and naïve hNPCs. Nine of the ten analytes including IL-1β, IL-6, TNF-α, IL-8, MMP-1, MMP3, MMP-12, NGF-β and VEGF-a in the ZNF865-VPR showed significant decreases in concentration compared to the non-target control, with the non-significant difference in MCP-1 trending towards a reduced level (**Figure 4J**). From the Luminex assay, a composite normalized SASP score was created that weighs each analyte equally and is normalized to the hNPC supernatants. Here, we saw significant differences in SASP score between the ZNF865-VPR dhNPCs and non-target control dhNPCs, as well as significant differences between the non-target control dhNPCs and hNPCs. The ZNF865-VPR dhNPCs had a SASP score that was slightly higher than the hNPCs, but this difference was not statistically significant (**Figure 4K**). This indicates that increased expression of ZNF865 leads to a decreased SASP profile and secretome more similar to hNPCs.

To assess ZNF865 regulation on ECM deposition within the disc, 3D pellet culture was performed using non-target dhNPCs pooled from three patient samples, and two ZNF865-VPR dhNPC patient samples (**Figure 4L**). After 3 weeks of culture, we saw significant increases in collagen production in both ZNF865-VPR patients compared to the pooled non-target pellets, with no significant increases in glycosaminoglycan (GAG) production (**Figure 4M,4N**). From these measurements, a composite anabolic score was created, weighing both collagen production and GAG production equally and normalizing to the non-target samples. Here we saw significant differences in anabolic score for both ZNF865-VPR samples compared to the non-target (**Figure 4O**). Additionally, when plotting the SASP scores and anabolic scores from both patients on the same plot we see a clear separation between the non-target samples and the ZNF865-VPR samples (**Figure 4P**), indicating that ZNF865 upregulation results in decreased inflammation and increased ECM deposition capability.

### ZNF865 overexpression significantly restores chromatin accessibility toward a healthy-like state

Given the close relationship between senescence and changes in chromatin landscape^44^, we performed Assay for Transposase-Accessible Chromatin using sequencing (ATAC-seq), with primary patient samples (**Table S5**), to identify differences in the chromatin accessibility associated with ZNF865 expression. When comparing hNPCs with dhNPCs we found 1163 sites that exhibit differential chromatin accessibility that were associated with 1705 genes identified by Genomic Regions Enrichment of Annotations Tool (GREAT)^45^. Further analysis of these genes revealed functional enrichment of 45 pathways, including glycosaminoglycan synthesis, MAPK signaling pathway, and cellular senescence (**Figure 5A, Table S6**). Upon upregulation of ZNF865 in dhNPCs, a total of 575 differential chromatin accessibility sites were identified, with 4 significantly enriched pathways compared to the dhNPCs (**Figure 5B, Table S7**). However, when comparing ZNF865-VPR dhNPCs to the hNPCs, we observed only 70 sites with differential chromatin accessibility (94% reduction) and no enrichment of any pathways (**Figure 5C, Table S8**). Principal component analysis using this cohort (**Figure 5D**) showed that dhNPCs become more related to hNPCs upon CRISPRa of ZNF865, suggesting that upregulation restores some molecular features of hNPCs. Similarly, additional RNA-seq was performed using comparable groups (**Table S9**), which showed a corresponding shift in gene expression when ZNF865 is upregulated in dhNPCs towards a healthier gene expression state (**Figure 5E**).

**Figure 5:**
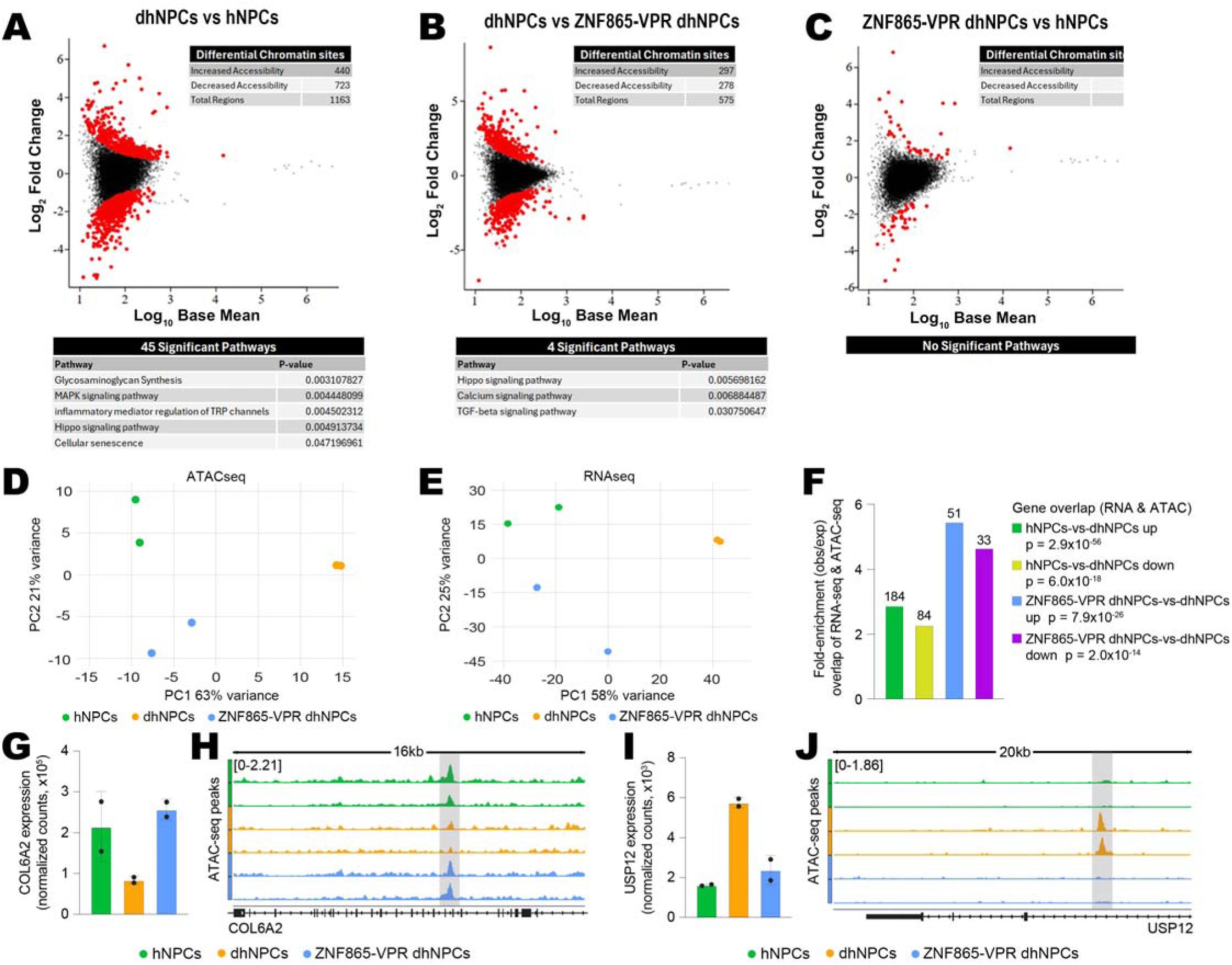
CRISPRa of ZNF865 returns human NP cells to healthy chromatin state and gene expression. **A** MA plots of the ATAC-seq data illustrating sites of differential chromatin accessibility comparing dhNPCs versus hNPCs, **B** dhNPCs versus ZNF865-VPR dhNPCs and **C** ZNF865-VPR dhNPCs versus hNPCs with relevant enriched pathways. **D** Principal component analysis of ATAC-seq data shows the relationship between hNPCs, dhNPCs, and ZNF865-VPR dhNPCs. **E** Principal component analysis of RNA-seq data shows the relationship between hNPCs, dhNPCs, and ZNF865-VPR dhNPCs. **F** Hypergeometric test results are shown as a bar plot to quantify the degree of gene-level overlap of RNA-seq and ATAC-seq datasets between comparisons of hNPCs and dhNPCs (green and yellow bars) and ZNF865-VPR dhNPCs and dhNPCs (blue and purple bars). Bars corresponding to the number of shared upregulated genes from RNA-seq and genes associated with increased chromatin accessibility from ATAC-seq for each comparison are denoted as “up.” Conversely, bars corresponding to the number of shared downregulated genes from RNA-seq and genes associated with decreased chromatin accessibility from ATAC-seq for each comparison are denoted as “down.” **G** Bar plot illustrates the differences in *COL6A2* expression between hNPCs dhNPCs, and ZNF865 VPR dhNPCs using normalized counts from RNA- seq. **H** Browser tracks display differences in ATAC-seq peaks (shaded region) in *COL6A2* from hNPCs, dhNPCs, and ZNF865-VPR dhNPCs. **I** Bar plot illustrates the differences in *USP12* expression between hNPCs, dhNPCs, a_2_n_8_d_7_ZNF865-VPR dhNPCs using normalized counts from RNA-seq. **J** Browser tracks display differences in ATAC-seq peaks (shaded region) in *USP12* from hNPCs, dhNPCs, and ZNF865-VPR dhNPCs.

To assess concordance between differentially expressed genes identified by RNA-seq and genes associated with changes in chromatin accessibility by ATAC-seq, we employed hypergeometric tests to quantify the degree of overlap between either upregulated or downregulated genes and genes associated with either increased or decreased chromatin accessibility. This analysis revealed statistically significant enrichment in the number of shared genes compared to what would be expected by chance across all comparisons **(Figure 5F)**. Specifically, we found 184 upregulated/increased accessibility and 84 downregulated/decreased accessibility genes that are shared between comparisons of hNPCs and dhNPCs, representing >3-fold and >2-fold enrichment compared to the amount of overlap expected by chance, respectively. Similarly, we found 51 upregulated/increased accessibility and 33 downregulated/decreased accessibility genes that are shared between comparisons of ZNF865-VPR dhNPCs and dhNPCs, representing >6-fold and >5-fold enrichment over the amount of overlap expected by chance, respectively.

Within the individual genes, there are several notable examples, including *COL6A2* **(Figure 5G)**, which was recently implicated as a potential marker of DDD^46^. Normalized counts from RNA-seq show upregulation of *COL6A2* in both hNPCs and ZNF865-VPR dhNPCs, compared to dhNPCs. These differences in *COL6A2* expression correlate with changes in chromatin accessibility, where we observed more prominent ATAC-seq peaks in both hNPCs and ZNF865-VPR dhNPCs compared to dhNPCs **(Figure 5H)**. We also found that *USP12*, which has a known functional role as an inflammatory mediator involved in MMP production^47^, is downregulated in both hNPCs and ZNF865-VPR dhNPCs compared to dhNPCs **(Figure 5I)**. These differences in expression correlate with changes in chromatin accessibility, where both hNPCs and ZNF865-VPR dhNPCs show less prominent ATAC-seq peaks compared to dhNPCs **(Figure 5J)**. Collectively, these results provide evidence that ZNF865 upregulation shifts dhNPCs toward a healthier molecular phenotype.

### CRISPRi of ZNF865 induces DDD and painful outcomes *in vivo*

To confirm that the associations observed between ZNF865 in the human patient samples, both in its relationship to age and back pain, and the molecular changes observed with ZNF865 regulation in human patient samples can directly drive pain outcomes, we knocked ZNF865 down directly in the rat interverbal disc with CRISPRi (**Figure 6A**). Prior to *in vivo* delivery, a rat orthologue of the human ZNF865 had to be verified. To begin, gRNA upstream of the ZNF865 orthologue from the Rat Genome Database^48^ were designed (**Table S10**), cloned into the dCas9-KRAB vector, and transduced into rat NP cells isolated from the caudal spine of Sprague Dawley rats before performing qPCR. Fold change in ZNF865 expression showed a significant reduction in gene expression in ZNF865-KRAB samples (**Figure 6B**). A significant reduction in cell proliferation was also observed (**Figure 6C**) alongside increases in the number of senescent cells via SA-β-gal and p16 staining (**Figure 6D, 6E, 6G**). Similarly to human samples, this was accompanied by an increase in DNA damage present within the cells (**Figure 6F**). Examining the expression of SASP related proteins showed increases in the secretion of MCP-1, and VEGF-A in the ZNF865-KRAB samples compared to non-target with ICAM-1 not reaching significance (*p*=0.0625). (**Figure 6H-6J**). Overall, the rat disc cells responded similarly to the human IVD cells with CRISPRi ZNF865 regulation.

**Figure 6:**
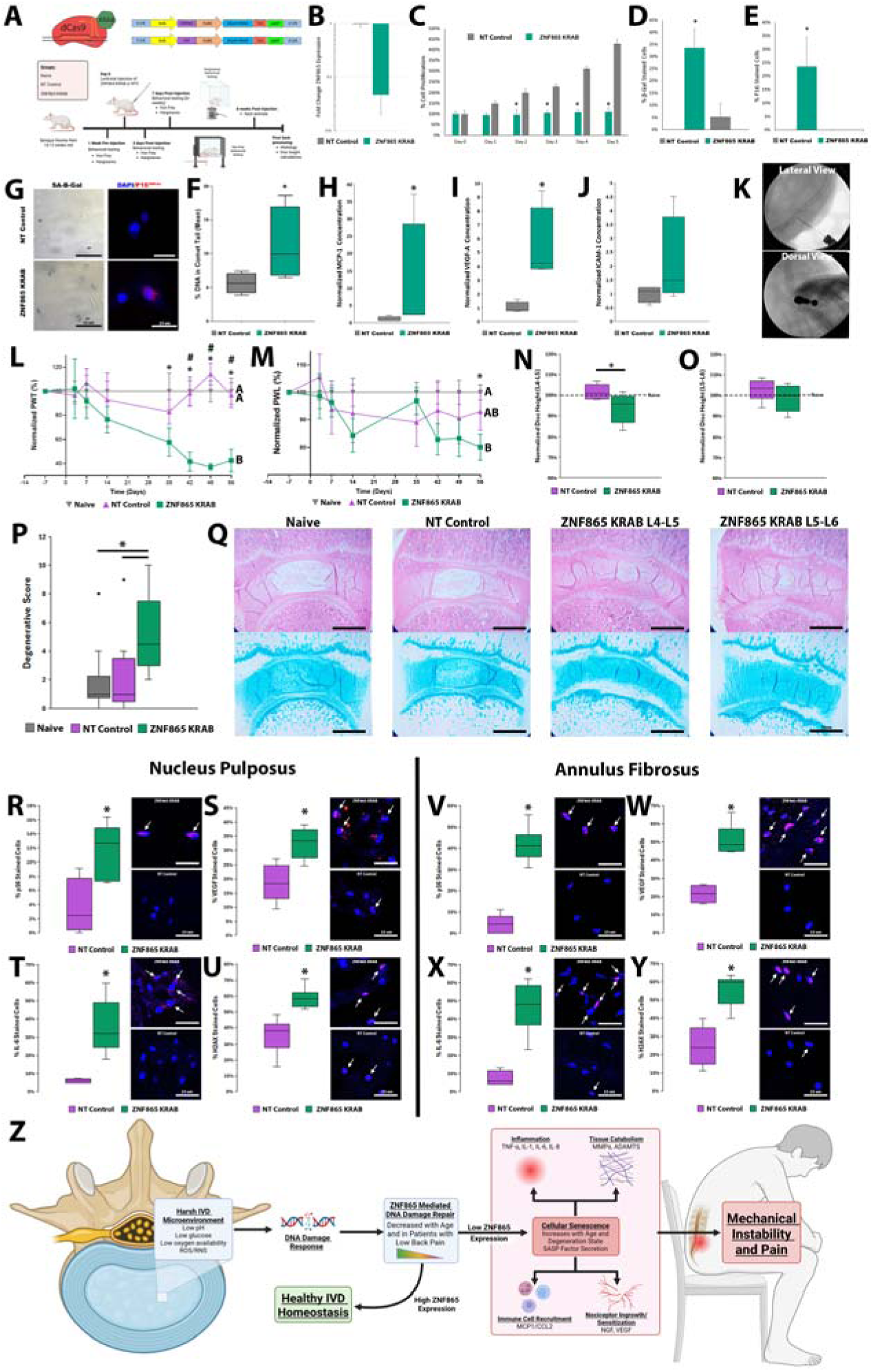
CRISPRi of ZNF865 induces DDD pathology and increases painful outcomes in vivo. **A** Experimental design of the 8-week animal model using the dCas9-KRAB repression system with plasmid maps for CRISPRi of ZNF865 or non-target. **B** Log fold change in ZNF865 expression **C** Quantified cell proliferation of rNPCs after transduction with ZNF865-KRAB or NT control vectors **D** Percent of SA-P-gal and **E** p16 stained cells 3 weeks after transduction (n=5). **F** Mean percent DNA in comet tail from alkaline comet assays (n=4). **G** Representative images from SA-β-gal, p16 and comet assays. **H** Normalized MCP-1, **I** VEGF-A, **J** and ICAM-1 concentrations. **K** Representative images of the fluoroscopy assisted lentiviral injections to the l4-L5 and L5-L6 IVDs. **L** Normalized paw withdrawal threshold (PWT) and **M** normalized paw withdrawal latency (PWL) (n=5-8). **N** Normalized disc height from the L4-L5 and **O** L5-L6 IVDs **P** Degenerative scoring and **Q** representative H&E and alcian blue staining for naive, NT-control and ZNF865 KRAB animals. **R** Percent nucleus pulposus cells stained for p16, **S** VEGF, **T** IL-6, and **U** H2AX with representative images. **V** Percent annulus fibrosis cells stained for p16, **W** VEGF, **X** IL-6, and **Y** H2AX with representative images (Nuclei=blue, protein=red, n=5-6). **Z** Proposed model of ZNF865 Function in IVD aging and degeneration. *(significance *=p<0.05).*

Following validation of the ZNF865 orthologue, ZNF865-KRAB or NT-control, were injected into the L4-L5 or L5-L6 discs of Sprague Dawley rats (**Figure 6K**), following protocols previously developed in our lab for targeted CRISPRi regulation of genes in the rat IVD^49^. Following injections, animals were evaluated for symptoms of mechanical and thermal nociception via von Frey and Hargreaves assessments respectively. Groups marked with different letters are significantly different (p<0.05). After 5 weeks a significant increase in mechanical pain sensitivity was seen in the ZNF865-KRAB group compared to both naïve and NT-control groups (**Figure 6L**). Thermal pain sensitivity also increased in the ZNF865-KRAB group compared to the naïve group at 8 weeks (**Figure 6M**). Following sacrifice, disc height was evaluated at each injected level. Here, a significant decrease in disc height was seen at the L4-L5 level but not the L5-L6 level between the ZNF865-KRAB group and the NT-control group (**Figure 6N, 6O**). Following staining with H&E and alcian blue, samples were scored for degeneration. ZNF865-KRAB groups showed significantly increased degenerative scores compared to both naïve and NT- control groups (**Figure 6P, 6Q**). Here, immunofluorescence staining also showed significant increases in p16, VEGF, IL-6 and H2AX staining in both the NP and AF regions of ZNF865 knockdown discs (**Figure 6R-6Y**). Additionally, decreases in paw withdrawal threshold are significantly correlated with increases in H2AX and p16 in both NP and AF regions, and VEGF in the AF (Table 1, Figure S4).

## Discussion

Despite the broad expression of ZNF865 across all cell types^50^, this study represents the first functional characterization of the ZNF865 gene. Through CRISPRa-mediated upregulation in three distinct cell types; *HEK293* cells, *ASC*s, and human NP cells, we identified a set of consistently enriched processes involving cellular senescence, DNA repair mechanisms, and cell cycle regulation pathways. These shared processes suggest that ZNF865 may act as a core regulator of molecular processes that underlie cell senescence and genomic stability.

We demonstrated that ZNF865 expression decreases with age and pathology, and that this reduction is associated with increased levels of DNA damage and cell senescence. Here we show that patient derived dhNPCs had significantly lower ZNF865 gene expression, lower protein levels, and greater DNA damage compared to hNPC controls, highlighting a potential connection between ZNF865 reduction and DDD in humans. This data is supported by single cell RNA sequencing of the caudal NP of aging mice *in vivo,* which showed a parallel decline in ZFP865 expression and the proportion of cells expressing ZFP865 with increased age. Additionally *in* vivo, we show that depletion of ZNF865 in a rat model is sufficient to induce histological degenerative outcomes and behavioral pain. Together, this data suggests that the loss of ZNF865 may represent a feature of disc tissue aging, contributing to aging related phenotypes and degeneration such as cell senescence which contributes to the lack of regenerative capacity of the disc.

CRISPR-based gene modulation systems such as CRISPRa and CRISPRi offer a method for modulating the expression of ZNF865, allowing us to define its functional roles. Using these CRISPR based gene modulation systems revealed that expression of ZNF865 is highly tunable and can lead to robust functional differences in both human and rodent models^32^, with gene expression shown to be associated with proliferation and cell cycle progression. Downregulation of ZNF865 in hNPCs led to reduced proliferation, increased senescence, and elevated DNA damage, all of which are features commonly associated with tissue degeneration and DDD. In rNPCs, these features were conserved, also showing reduced proliferation and increases in senescence, DNA damage and in SASP protein production including MCP-1 and VEGF-A. Conversely, when ZNF865 was upregulated in dhNPCs, these phenotypes were reversed, with increased proliferation rates, decreased senescence marker staining, and decreased DNA damage. This also led to a significant reduction in many SASP factors such as proinflammatory cytokines (IL-1, IL-6, TNF-α, IL-8), proteases (MMP-1, MMP-3, MMP-12), and pain related growth factors (NGF, VEGF) commonly associated with discogenic back pain. After upregulation of ZNF865, these dhNPCs showed cell phenotypes more similar to the hNPCs. Additionally, in 3D pellet culture ZNF865 upregulation showed increased ECM deposition capability.

These phenotypic differences between patient samples are consistent with gene regulatory changes. Both RNA-seq and ATAC-seq showed that CRISPRa of ZNF865 recapitulated global features of gene expression and chromatin accessibility found in hNPCs. This effect can be seen at the level of individual genes, where we observed many up- and down-regulated genes that also harbored sites of more open and closed chromatin. One notable example we identified is *COL6A2,* whose expression is higher in hNPCs and ZNF865-VPR dhNPCs compared to dhNPCs. This gene encodes a type VI collagen chain, revealing a potential genetic link with increased ECM deposition by ZNF865-VPR dhNPCs in 3D pellet culture. Overall, these data highlight concordant changes in phenotypes and gene regulation, where our data suggests ZNF865 drives an overall shift of both gene expression and chromatin state toward that of healthy NP cells.

*In vivo,* we demonstrate that intradiscal CRISPRi knockdown of ZNF865 induces behavioral pain responses. Our behavioral data showed a significant increase in mechanical pain sensitization, and to a lesser degree, thermal pain sensitization in the ZNF865 knockdown group. These behavioral outcomes parallel the nociceptive changes observed in well-established models of disc degeneration^49,51^ and support the role of ZNF865 in regulating pain-relevant pathways within the IVD. This was further corroborated by histological analysis in which ZNF865-KRAB injected discs exhibited reduced disc height at the L4-L5 level and increased degenerative scores across both levels, with this difference perhaps due to differences in load mechanics. Despite differences, the induction of pain in the absence of a mechanical injury challenges traditional models of discogenic pain. Most rodent disc degeneration models rely on large needle puncture, enzymatic digestion, or biomechanical overloads, all of which introduce acute structural damage to the disc, which may confound interpretation of pain mechanisms. Here nociceptive sensitization was achieved through transcriptional downregulation, which highlights molecular dysfunction as a primary driver of pain rather than structural disruptions. Additionally, the gradual progression of pain behavior seen in the von Frey assessment mirrors the slow progression of degeneration seen in human disc disease, suggesting that ZNF865 knockdown initiates a biological cascade that cumulatively leads to degeneration and painful symptoms, more similar to human pathology. This pain behavior was also strongly correlated (**Table 1**) to increases in DNA damage markers, senescence markers, and pain related growth factor secretion, supporting proposed mechanisms (Figure 6Z) of ZNF865 induced discogenic pain development.

Collectively, this data suggests that ZNF865 plays a fundamental role in DDD and aging, specifically in maintaining genetic stability through DNA repair mechanisms that prevent the onset of senescence (**Figure 6Z**). The rescue of dhNPCs from a senescent phenotype upon ZNF865 upregulation not only points to the importance of ZNF865 within the degenerative phenotype but also identifies ZNF865 as a potential therapeutic target for DDD. Previously, we demonstrated upregulation of ZNF865 in multiplex with ACAN and Col2a1 enhanced functional tissue deposition and improved biomechanical properties of human-scale engineered discs^32^. This suggests that in addition to decreasing inflammatory pathways caused by DNA damage and SASP, CRISPR gene therapies using ZNF865 have potential to be used in multiplex for a wide variety of purposes including increased protein production or mechanical properties of engineered tissues.

This work also offers broader implications for other aging related degenerative diseases, where controlling cellular senescence and SASP may be paramount to understanding and slowing disease progression. Ultimately, preventing senescence may improve the health span of individuals by preventing the onset of many musculoskeletal or neurodegenerative disorders. ZNF865 was recently identified independently in a family of zinc finger proteins where increased gene expression was associated with better prognosis in esophageal cancer^52^, indicating that it may be a useful clinical target for understanding cancer progression. ZNF865 was also identified as a potentially relevant target in Parkinson’s disease^53^. *Liu et al.* suggested that a microRNA associated with ZNF865 protein expression can alleviate damage caused by Parkinson’s disease by preventing apoptosis and enhancing motor neuron performance^53^. Within our RNA sequencing data, several neurodegenerative diseases such as ALS, Parkinson’s and Alzheimer’s were listed as top pathways effected by upregulation of ZNF865, suggesting that targeting ZNF865 or its downstream pathways could represent a strategy for restoring tissue function in aging related degenerative diseases more broadly however this requires further experimental validation.

This work indicates ZNF865 is an important gene for healthy cell function and genome maintenance. However, key questions remain that need to be clarified in future experiments. Identification of the upstream regulators and downstream effectors through DNA binding studies (ChIPseq) will allow for better regulatory networks to be established around ZNF865 and help to define the molecular interactions leading to the observed results. However, current antibodies are insufficient to perform such studies and FLAG-tag ZNF865 constructs have been built and are a key piece of future work around this gene/protein. Additionally, in general, novel tools need to be developed to help study this gene/protein, including the development of blocking antibodies, inhibitors or activators that could allow for enhanced control of ZNF865 regulation outside of using CRISPR-based therapies. This work uses DDD as a model system for understanding DNA damage and senescence regulation by ZNF865 using the highest knockdown gRNA target, however, additional knockdown levels, timepoints, primary cell types and disease models should be considered to establish a broader understanding of the functional role that ZNF865 plays within aging related diseases. This is especially important considering the rare genetic disorders that have been associated recently with ZNF865 mutations^38^. Additionally, healthy patient samples used in this study are obtained from trauma patients, but there may be asymptomatic degeneration present in these patients that is not accounted for, which will require a larger clinical investigation in the future. This data set strongly supports larger future clinical studies looking at larger patient populations and co-morbidities to fully understand the relationships observed here in the broader patient population.

Overall, this work also highlights the use of CRISPR gene modulation strategies for probing unknown gene function and identifies ZNF865 as an important gene in regulating cell cycle and DNA repair mechanisms, with significant implications for DDD related pain and other senescence associated degenerative disorders. This study serves as a first step in understanding the basic biology of the ZNF865 gene and establishes the importance of understanding ZNF865 dysregulation for both disease pathology, and as a potential therapeutic target for DNA damage/senescence associated disorders.

## Acknowledgements

Research reported in this publication was supported by the National Institute of Health under award numbers R01 AR083990, R01 AR074998 and R01 AR080096. Sequencing was performed at the DNA Sequencing Core Facility, University of Utah. FACs was performed at the Flow Cytometry Core Facility, University of Utah. Research reported in this publication utilized the High-Throughput Genomics Shared Resource at the University of Utah and was supported by NIH/National Cancer Institute (NCI) award P30 CA042014. The content is solely the responsibility of the authors and does not necessarily represent the official views of the NIH.

## Author Contributions

Conceptualization: CL, HL, JH, JTP, TDJ, MC, SEG, BD, JG, RDB

Methodology: CL, HL, JH, JTP, BD, JG, RDB

Investigation: CL, HL, JH, JTP, TDJ, JM, MC

Visualization: CL, HL, JH, JTP, TDJ, MC, RDB

Funding acquisition: JCI, RDB

Project administration: CL, RDB

Supervision: JCI, JG, RDB

Writing-original draft: CL, HL, JH, JTP, TDJ, RDB

Writing-review and editing: CL, HL, JH, JTP, TDJ, MC, SEG, BD, JCI, JG, BL, RDB

**Competing interests:**

Authors declare that they have no competing interests

**Data and materials availability:**

GEO accession # GSE311657

## Materials and Methods

### Experimental overview

CRISPRa was used to upregulate the expression of ZNF865 in HEK 293 cells, ASCs and human dhNPCs and investigated for changes in gene expression via RNAseq. Next, ZNF865 expression and DNA damage were examined in healthy asymptomatic NPCs and dhNPCs to investigate DDD pathology. An aging mouse model from *Jacobsen et al.* was used to investigate an identified orthologue for ZNF865 expression changes with aging^54^. CRISPRi was then used to downregulate the expression of ZNF865 in hNPCs to investigate differences in proliferation, senescence and DNA damage as a result of decreased expression. Next, CRISPRa was used to upregulate the expression of ZNF865 in dhNPCs to investigate differences in proliferation, senescence, SASP expression, DNA damage and chromatin accessibility as a result of increased expression. Finally, a rat orthologue of ZNF865 was identified and validated *in vitro* through CRISPRi of ZNF865 in rat NP cells, and an *in vivo* model run examining behavioral pain responses and degenerative outcomes.

### General cell culture

All cell culture was performed in standard culture conditions (21% O_2_, 5% CO_2_, 37°C), with media changes every 2-3 days. *HEK 293*: Complete growth medium for cell culture for HEK 293 (ATCC CRL-1573) cells consists of HG-DMEM (Thermo) supplemented with 10% fetal bovine serum (FBS) (Thermo), 25μg/mL gentamicin (Corning), and 25mM HEPES (Thermo). *hASC:* Complete growth medium for culturing of human adipose derived stem cells (ASCs, ATCC SCRC-4000) consists of Lonza ADSC Basal Medium (Lonza, Lexington, MA PT-3273), 10% MSC FBS (Thermo), 5mL GlutaMax (Thermo), 30μg/mL Gentamicin (Corning), and 15ng/mL Amphotericin (Thermo). *hNPCs:* Primary human nucleus pulposus cells (hNPCs) were cultured in complete growth medium containing DMEM-HG (Thermo) supplemented with 10% FBS (Thermo), 50μg/mL gentamicin (Corning), 25mM HEPES (Thermo), and 2ng/mL of recombinant human fibroblast growth factor-basic (rhFGF, Peprotech). *rNPCs:* Primary rat nucleus pulposus cells (rNPCs) were cultured in Hams-F-12 Culture Media (Thermo), supplemented with 20% FBS (Thermo), 25ug/mL gentamicin (Corning), 8 mM HEPES, and 1.25 ug/mL amphotericin b (Thermo). Cells were cultured to 90% confluency before passaging 1:3 with 0.25% trypsin/EDTA solution (Thermo) as previously described^55^.

### gRNA design

Upregulation gRNA design was performed using Genome Target Scan 2 and ChopChop^56,57^. The 5’-UTR, the promoter region up to 500bp upstream of the human ZNF865 transcription start site (TSS), and up to 500bp upstream of the second exon were analyzed for gRNAs. The top 6 gRNAs from each analysis tool, with the least number of off-target sites, were compared and inspected using the BLAT tool of the UCSC genome browser to ensure minimal overlap between gRNAs and optimal placement near DNase hypersensitive regions (**Table S10**)^58^. A total of 6 gRNAS were selected for validation *in vitro*, as well as a gRNA that does not target the human genome (non-target control/NTC). Oligos for ZNF865 gRNAs were synthesized by the University of Utah DNA/Peptide Synthesis core and Thermo Fisher Scientific.

Downregulation gRNA design was performed using Genome Target Scan 2 (GT-Scan2) and ChopChop^56,57^. Briefly, a panel of 5 gRNAs were designed within the 5’-UTR and the promoter region up to 1000 bp upstream of the human ZNF865 TSS (**Table S10**). Oligos for ZNF865 gRNAs were synthesized by the University of Utah DNA/Peptide Synthesis core and ThermoFisher Scientific.

Downregulation gRNA design in the rat genome was performed using Genome Target Scan 2 (GT-Scan2) and ChopChop^56,57^. The rodent ZNF865 sequence was identified using the Rat Genome Database (RGSC Genome assembly v6.0)^48^. Here, the 5’UTR, the promoter region up to 1000 base pairs upstream of the identified rat ZNF865 transcription start site were analyzed for gRNAs. The top 15 gRNAs with the least number of off-target sites, from each analysis tool were inspected using the BLAT tool in the UCSC genome browser to ensure minimal overlap between gRNAs (**Table S11**). These 15 gRNA were then selected for validation as well as a gRNA that does not target the rat genome (non-target/NT). Oligos for ZNF865 were synthesized by ThermoFisher Scientific.

### Cloning

gRNAs, and a non-target that does not target the human or rat genome, were synthesized, annealed, phosphorylated, and ligated into individual U6-gRNA-UbC-DsRed-P2A-Bsr (upregulation gRNA) lentiviral upregulation expression vectors (Addgene, #83919) or KRAB CRISPRi downregulation expression vector pLV-hUbC-dCas9-KRAB-T2A-GFP (Addgene, #71237). Successful gRNA insertion was verified through Sanger sequencing^55,59,60^.

### Harvesting of primary hNPCs and culture

Human healthy and degenerative nucleus pulposus tissues were obtained from surgical tissue waste from trauma patients with no history of back pain, and patients undergoing surgical treatment for back pain respectively (Category 4 exemption verified by University of Utah IRB). Briefly, NP tissue was rinsed twice with washing medium (DMEM-HG, 165 μg/mL gentamicin sulfate, 100μg/mL kanamycin sulfate, 1.25μg/mL amphotericin), minced, and enzymatically digested in washing medium with 0.3% (w/v) collagenase type II (Worthington Biochemical), 0.2% pronase (Sigma), and 5% FBS (Thermo) for 2-3 hours at 37°C with 5% CO_2_ under gentle agitation ^55^.

Isolated cells were passed through a 70μm cell strainer and washed twice. Cells were counted and plated at a density of 10,000 cells/cm^2^ in NP cell culture medium (DMEM-HG with 10% FBS, 50μg/mL gentamicin, 25mM HEPES) supplemented with fresh 2ng/mL fibroblast growth factor-2 (FGF-2) (Peprotech). Cells were cultured in this medium at 37°C and 5% CO_2_ in a humidified atmosphere and subcultured to 90% confluency, as previously described^55^.

### Harvesting of primary rat NPCs and culture

Rat IVD tissue was obtained from the caudal spine (tails) of sacked Sprague Dawley rats. Upon acquisition, tails were macroscopically dissected to remove skin and connective tissue surrounding the spine. Once IVD were located AF and NP tissue were separated and placed in washing medium (DMEM-HG, 165 ug/mL gentamicin sulfate, 100ug/mL kanamycin sulfate, 1.25ug/mL amphotericin b). NP tissue was then digested in 0.01% pronase (Millipore Sigma), 0.0125% collagenase type 2 (Worthington Biochemical) and 5% FBS (Thermo) overnight at 37°C, Following incubation, cells were centrifuged and washed with warm PBS (Thermo) twice before being passed through a 70 um cell strainer. Cells were then re-centrifuged and re-suspended in culture media and plated in flasks pre-coated with LN5 supernatant and cultured until use at passage 1-2.

### Lentivirus production

The gRNA plasmid DNA was used to produce lentivirus, as previously described^55,59,60^. The amplified gRNA plasmid DNA was co-transfected into HEK 293T cells with psPAX2 (Addgene, plasmid #12260) and pMD2.G (Addgene, plasmid #12259) lentiviral packaging plasmids to create a lentivirus, as previously described^45^, Briefly, HEK 293T cells were seeded at a density of 62,500 cells/cm^2^. The following day, lentiviral plasmids were added to the cells with Lipofectamine 2000 (Thermo), following the manufacturer’s protocol. After 24 hours, the cell supernatant was discarded and replaced with fresh medium. Cell supernatant containing the lentiviral vectors was collected at 48 and 72 hours, concentrated to 100X, and stored at -80°C until use^60^.

### dCas9-VPR upregulation transduction

*HEK 293/ASC:* HEK 293 cells and ASCs were plated at a density of 20,000 cells/cm^2^ and 5,000 cells/cm^2^, respectively, allowed to attach overnight, and transduced with EF-1α-dCas9-VPR-PuroR (dCas9-VPR) (Addgene, #99373) lentivirus, (1:20 dilution) in growth medium containing 4μg/mL polybrene, the following day. Both cell types were subjected to antibiotic selection (1μg/mL) over the course of 3 days to select for dCas9-VPR expressing cells. *hNPCs:* hNPCs (p1-p2) were plated at a density of 20,000 cells/cm^2^, allowed to attach overnight, and transduced with EF-1a-dCas9-VPR-eGFP (Addgene, #241307) lentivirus, (1:2 dilution) in growth medium containing 4μg/mL polybrene, the following day. 72-hours following transduction hNPCs were subjected to fluorescence activated cells sorting (FACS) to select for GFP+ expressing cells.

After dCas9-VPR transduction, dCas9-VPR expressing HEK 293 cells, hNPCs, and ASCs were transduced with gRNA targeting lentiviral vectors. *HEK293-VPR:* HEK 293-VPR cells were plated at a density of 20,000 cells/cm^2^ and transduced with gRNA vector (1:250 dilution) targeting ZNF865 (Addgene, #241309) or non-target the following day in complete media. *hNPC-VPR*: hNPCs were plated at a density of 5,000 cells/cm^2^ in a 24-well plate and transduced with gRNA vector (1:160 dilution) targeting ZNF865 (Addgene, #241309) or non-target in complete media the following day. *ASC-VPR*: ASCs were plated at a density of 5,000 cells/cm^2^ in a 24-well plate and transduced with gRNA vectors (1:160 dilution) targeting ZNF865 (Addgene, #241309), or non-target in complete media the following day. Transduced HEK 293, ASCs and hNPCs were examined for fluorescence after 72 hours and showed near 100% transduction efficiency.

### dCas9-KRAB downregulation transduction

For ZNF865 downregulation, naïve HEK-293, naïve ASCs, naïve hNPCs, and naïve rNPCs were plated at a density of 20,000 cells/cm^2^, 5,000 cells/cm^2^, 10,000 cells/cm^2^ or 10,000 cells/cm^2^ respectively, in 24-well plates and allowed to attach overnight. The following day, 100X sgRNA targeting ZNF865 in the human or rat (Addgene, #241308) or non-target virus was diluted 1:16 in complete growth medium supplemented with 4μg/mL polybrene and used to transduce cells. Cells were examined for fluorescence after 72-hours and showed near 100% transduction efficiency.

#### qRT-PCR

Successfully transduced cells were analyzed for changes in ZNF865 gene expression by qRT-PCR (n=4). 72-hours post-transduction, RNA was isolated and harvested using the Quick-RNA Micro Kit (Zymo Research, R1051), complementary DNA (cDNA) was synthesized from the purified RNA with high-capacity cDNA reverse transcription kit with RNAse inhibitor (Applied Biosystems). cDNA was then used for qRT-PCR with TaqMan gene expression assays (Thermo) for human ZNF865 (Hs05052648_s1) or rat ZNF865 (Custom Assay ID APT2HZ6). Beta-2-microglobulin (B2M, Hs00187842_m1) was used as an internal standard and changes in ZNF865 expression was normalized to B2M expression^61^. Fold-change in mRNA expression relative to the non-target or ZNF865 edited cells was calculated using the ΔΔCT method.

### RNA-sequencing

RNA-sequencing (RNAseq) was utilized to evaluate differential gene expression due to ZNF865 upregulation. HEK 293 cells, ASCs, and primary human NP cell samples (**Table S7**) were analyzed for global changes in gene expression after modification with ZNF865 or NTC CRISPRa vectors. Briefly, Total RNA was isolated from samples using a Quick-RNA Kit (Zymo Research, Irvine, CA). Isolated Total RNA (100-1000ng) was prepared using an Illumina TruSeq Stranded mRNA Library Prep Kit with PolyA Selection and samples were submitted to the High-Throughput Genomics Shared Resource Core at the Huntsman Cancer Institute for sequencing on a NovaSeq 6000, utilizing a NovaSeq Reagent Kit v1.5_150x150bp sequencing with 33 million reads per sample.

Using previously described methods, data were normalized and compared to non-target control cells^55,59^. Sequencing reads were aligned to the hg38 build of the human genome using HISAT2 ^62^and SAMtools ^63^ was used to convert SAM files to BAM files. Genes were defined by the University of California Santa Cruz (UCSC) Known Genes, and reads mapping to known genes were assigned using featureCounts from the SubRead package^64,65^. Reads were normalized and differential analysis was performed using EdgeR ^66^ (**Figure 1E-1G**) or the DESeq2 package in R (**Figure 5E**)^64,67^. We used an adjusted p-value threshold of <0.05 to identify genes that were significantly up- or down-regulated when comparing patient samples in Figure 6. Enriched GO biological and molecular processes were determined from lists of significantly differentially expressed genes using Enrichr ^39,68^.

### Single cell RNA-sequencing

Single cell RNA sequencing (scRNA-seq) previously performed in *Jacobsen et al.* was mined for the relationship of ZFP865, the identified orthologue of human ZNF865^41^, with aging in mice. In this study, mice were used at three ages: 4 months (skeletal maturity), 12 months (equivalent to human age with peak back pain disability), and 24 months (equivalent to human age with peak back pain prevalence)^54^. Cells were isolated from 6 whole naïve coccygeal IVDs per mouse, from 6 mice per age group and pooled. Samples were processed by the Genomics Core Facility at the Icahn School of Medicine at Mount Sinai. Samples were processed using the 10X Genomics Chromium 3’ Kit and sequenced using an Illumina S1 NovaSeq chip. Cell Ranger software mapped reads to mouse mm10-2020-A reference genome. Data was processed via Seurat including quality control filtering, normalization, and unsupervised clustering. Cell clusters were visualized with uniform manifold approximation and projection (UMAP). Differential gene expression analysis identified gene markers for individual clusters and canonical markers facilitated annotation of clusters. Data was subset to NP and Notochordal clusters for analysis of ZFP865 expression. ZFP865 expression level was determined at the cellular level for each age, percentage of cells positively expressing ZFP865 at each age was also determined.

### Cell proliferation quantification

Following successful transduction, HEK 293, ASCs, and hNPCs transduced with either the CRISPRa upregulation system or CRISPRi downregulation system (n=4) were evaluated for cell proliferation over the course of 3-10 days. Individual cells were counted using ImageJ or a hemacytometer^69^. Briefly, fluorescing cell counts were obtained by uploading images into a stack and thresholding the images to ensure only individual cells are shown within the image. After thresholding, cell counts were obtained by analyzing particles, generating a mask, and then counting the masks generated while excluding cells on the edges of the image. In instances where cell density was too great for thresholding and visually isolating individual cells, cells were manually counted in ImageJ using the cell counter plugin^69^. For each primary patient sample, 3-5 technical replicates were run alongside each other to establish a patient average, reducing variability within each patient.

### Immunofluorescence (P16^INK4A^, p21^CIP^^1^ and ZNF865)

Primary hNPCs or rNPCs were plated into cell culture chamber slides at a concentration of 10,000 cells/cm^2^ and then transduced with the ZNF865 or NTC downregulation expression cassettes. Cells were then cultured for 10 or 24 days after which the cells were fixed with 10% formalin. Following fixation cells were permeabilized with 0.2% Triton X in PBS for 5 min. Cells were then washed three times with PBS and treated with blocking solution (5% goat serum [MP Biomedicals] in PBS for 1 hour at room temperature). Following treatment with blocking solution, the blocking solution was removed, and the cells were treated with primary antibody; p16^INK4A^ (p16) antibody (1:100 in 1% BSA in PBS; Proteintech), p21^CIP^^1^ (p21) antibody (1:100 in 1% BSA in PBS; Proteintech) or ZNF865 (1:100 in 1% BSA in PBS; Atlas Antibodies), normal rabbit IgG (Invitrogen) or BSA only negative controls and incubated at room temperature for 2 hours. Following incubation cells were washed three times with PBS, treated with goat anti-rabbit Coralite 594 (1:100 Proteintech) in 1% BSA in PBS, and incubated for 1 hour at room temperature. Subsequently, cells were rinsed three times with PBS, treated with DAPI solution (3ng/mL) for 10 minutes, and mounted with Vectamount Mounting Medium prior to fluorescence imaging (Olympus IX73). For each primary patient sample, 3-5 technical replicates were run alongside each other to establish a patient average, reducing variability within each patient.

### Alkaline comet assay

The alkaline comet assay was used to assess DNA damage in primary hNPCs and rNPCs. NP cells were trypsinized and resuspended in cold PBS at 1 x 10^6^ cells per mL. 5 x 10^4^ NP cells were mixed in a 3:7 volume ratio with 1% Low Melting Point Agarose (BP1360, Fisher Scientific). The mixture was then applied to a Superfrost Plus slide (12-550-15, Fisher Scientific) precoated with 1% normal melting point agarose (20-102, Apex). The slides were then transferred to the lysis solution (2.5 M NaCl, 100 mM disodium EDTA, 10 mM Tris Base, 1% Triton X, pH 10) and allowed to lyse overnight at 4°C in the dark. Immediately following lysis, the slides were washed 1x with dH_2_O and immersed in an electrophoresis tank containing alkaline electrophoresis solution (300 mM NaOH, 1 mM disodium EDTA, pH >13) for 30 min followed by electrophoresis at 1 V/cm for 20 min at 300 mA. Slides were then washed for 10 minutes in PBS and an additional 10 minutes in dH_2_O at 4°C and allowed to dry at RT overnight. Dried slides were then stained with 1ug/mL DAPI (62248, Thermo). At least 100 randomly selected stained nuclei per slide were imaged under an Olympus IX73 microscope. Comets were analyzed in ImageJ using the OpenComet plugin^70^.

### SA-β-Galactosidase staining

Primary hNPCs or rNPCs were plated in a 24-well plate at a concentration of 10,000 cells/cm^2^ and allowed to attach overnight. The following day attached cells were transduced with the ZNF865 (Guide 2 and Guide 3) or NTC downregulation expression cassettes. Cells were cultured for 10 or 24 days, at which point they were fixed and stained for SA-β-galactosidase using a Senescent Cell Histochemical Staining Kit (Sigma-Aldrich, CS0030-1KT). Cellular senescence was quantified by counting SA-β-galactosidase stained and unstained cells using ImageJ. For each primary patient sample, 3-5 technical replicates were run alongside each other to establish a patient average, reducing variability within each patient.

### Custom Human Luminex assay

Following upregulation of ZNF865 in primary dhNPCs a custom Luminex panel (Procartaplex 11 analyte multiplexed assay, Thermo) was run for IL1-β, IL-6, IL-8(CXCL8), TNF-α, MCP-1(CCL2), MMP-1, MMP-12, MMP-3, MMP-8, NGF-β, VEGF-a. 10-14 days following transduction of dhNPCs with ZNF865-VPR or NT, degenerative and asymptomatic samples were passaged and plated in T-25 flasks at the appropriate cell density and 3 mL of media added per flask. 3 days following, supernatant was collected and spun at 1400 rpm for 10 minutes to remove particulates, then aliquoted for storage at -80°C until use. Images were taken at the same time point as collection for normalization based on cell number. The Procartaplex plate was then run according to the manufacturer’s instructions using 50 µl of supernatant per sample per well and run on a Luminex Magpix. Each sample was run in duplicate. After running the plate, values were then averaged across each duplicate and divided by the cell density to account for differences in cell numbers. Values below the detection range were given concentrations of ½ the detection limit of each analyte. ZNF865-VPR dhNPC values were then normalized to the non-target across individual patients before averaging across all patients to get a normalized concentration for each analyte. A composite SASP score weighting each analyte equally was then calculated by normalizing each analyte to the highest concentration measured across all samples, then taking the sum across all analytes to get a composite score for each sample.

### Custom Rat Luminex assay

Following downregulation of ZNF865 in the rNPCs a custom Luminex panel (Procartaplex 9 analyte multiplexed assay, Thermo) was run for IL1-β, IFN-γ, IL-6, IL-10, TNF-α, MCP-1(CCL2), NGF-β, VEGF-a, and ICAM-1. 21 days following transduction of rNPCs with ZNF865-KRAB or NT, cells were passaged and plated in 6 well plates at the appropriate cell density and 1.5 mL of media added per flask. 3 days following, supernatant was collected and spun at 1400 rpm for 10 minutes to remove particulates, then aliquoted for storage at -80°C until use. Images were taken at the same time point as collection for normalization based on cell number. The Procartaplex plate was then run according to the manufacturer’s instructions using 50 µl of supernatant per sample per well and run on a Luminex Magpix. Each sample was run in duplicate. After running the plate, values were then averaged across each duplicate and divided by the cell density to account for differences in cell numbers.

### Pellet Culture of dhNPCs

To evaluate extracellular matrix (ECM) deposition in the ZNF865 edited cells, 3D pellet cultures were performed. ZNF865-VPR dhNPCs and non-target dhNPCs were passaged and resuspended at 1.25 million cells/mL in serum free growth medium. Serum free media consisted of DMEM-HG with pyruvate (Thermo), 0.1 µm dexamethasone, 1% antibiotic/antimycotic, 0.17mM ascorbic acid 2-phosphate, 0.35mM proline and 1% ITS+. Due to low cell numbers, three non-target dhNPC samples were pooled to form a single non-target dhNPC group. 200 µL aliquots of cell suspension was pipetted into individual wells of 96-well v-bottom plate and spun at 270G for 10 min at 4°C. Cells were allowed to contract for 48 hrs to form pellets. Pellets were then gently lifted from the bottom of the plate. Media was changed every 3 days for 21 days and supernatant collected during every media change.

### Pellet biochemical analysis

The total amount of collagen content within papain digested pellets and supernatant was analyzed using a modified hydroxyproline assay as previously described ^32,60^. The total amount of DNA and GAG content within papain digested pellets and supernatant was analyzed using a previously described Hoescht dye assay and dimethylmethylene blue (DMMB) assay, respectively ^32,60,71,72^. A composite anabolic score weighting the contributions of collagen and GAG equally by normalizing to the highest concentration measured across all samples, then taking the sum of each normalized collagen and GAG score to get a complete anabolic score. These were then normalized to the average non-target dhNPC value.

### ATAC-sequencing

Primary patient samples (**Table S4**) were prepared in technical duplicate, where 50,000 cells per replicate were taken from cryogenically frozen vials and centrifuged at 500 x g for 5 minutes at 4°C. The supernatant was aspirated, and the resulting cell pellet was washed with 250 µL of ice-cold 1X PBS, pH 7.4 (Thermo; 10010072). The cells were then centrifuged at 500 x g for 5 minutes at 4°C, the supernatant was aspirated, and the resulting cell pellet was resuspended in 50 µL of lysis buffer containing the following: 10 mM Tris-HCl, pH 7.4 (Sigma-Aldrich; T2194), 3 mM MgCl_2_ (Sigma-Aldrich; M1028), 10 mM NaCl (Sigma-Aldrich; 71386), 0.1% NP-40 (Millipore Sigma; 492016), 0.1% Tween-20 (Bio-Rad; 1610781), and 0.01% digitonin (Invitrogen; BN20061). Samples were then lysed via incubation on ice for 15 minutes and centrifuged at 500 x g for 5 minutes at 4°C. Following lysis, 1 mL of wash buffer (10 mM Tris-HCl, pH 7.4, 3 mM MgCl_2_, 10 mM NaCl, and 0.1% Tween-20) was added to each sample, which was then centrifuged at 500 x g for 5 minutes at 4°C. The supernatant was aspirated, and the resulting cell pellet was resuspended a second time in 1 mL of wash buffer. To pellet nuclei, cells were centrifuged at 500 x g for 10 minutes at 4°C. The supernatant was aspirated, and each resulting nuclei pellet was resuspended in transposition buffer containing the following: 10 mM Tris-HCl, pH 8.0, 5 mM MgCl_2_, 10% Dimethylformamide (Fisher Bioreagents; BP1160), and 1 µL of TDE1 Tagment DNA enzyme per sample (Illumina; 20034197) diluted in PBS, pH 7.4.

For the transposition reaction, nuclei were incubated at 37°C for 30 minutes in an Eppendorf ThermoMixer C (EP5382000015) set to rotate at 1000 RPM. Following transposition, DNA was purified using the Zymo DNA Clean & Concentrator-5 kit (D4014) according to the manufacturer’s instructions with a binding buffer to sample ratio of 5:1. The purified DNA was then amplified by PCR using a final concentration of 1X Phusion Master Mix (New England Biolabs; M0530), and 200 nM each of forward and reverse barcode primers. Amplified libraries were purified using a 1:1 volume ratio of AMPure XP beads (Beckman Coulter; A63882) and fragment size distribution and quality were assessed using a High-Sensitivity D1000 ScreenTape assay.

Libraries were sequenced on a NovaSeq X at a depth of ∼50 million reads/sample and 100 bases from the 3’ end of each read were trimmed using seqtk^73^ with the following parameters: trimfq -e 100. Trimmed reads were then aligned to the hg38 build of the human genome using Bowtie^74^ with the following parameters: --chunkmbs 512 -m 1 -t --best -q -S -l 32 -e 80 -n 2. SAM files were converted into BAM files and sorted using SAMtools^63^. Peaks representing regions of accessible chromatin were called using MACS2^75^ with an adjusted P-value cutoff of <0.05 and an mfold parameter between 15 and 100. FeatureCounts^65^ was used to quantify reads that aligned to ATAC-seq peaks called across all samples. Reads were normalized and chromatin regions displaying differential accessibility were identified using the DESeq2 package for R^67^. To identify regions of chromatin with differential accessibility, we compared dhNPC samples to both hNPCs and ZNF865-VPR dhNPCs, and ZNF865-VPR dhNPCs to hNPCs.

### Animals

All procedures were performed with the approval of the University of Utah Institutional Animal Care and Use Committee (IACUC). Here, 24 Male Sprague Dawley rats 10-12 weeks of age were used. Animals were allowed free movement in their home cages and were dual housed throughout the course of the study. Rats were allowed food and water *ad libitum* throughout. Routine health and wellness exams were performed and wight were performed weekly to assess general animal health. Rats were maintained at 72°F-74°F and 12/12h light/dark cycles (6am-6pm lights on).

### Non-Invasive, Image Guided. Cannulated Lentiviral Injections

Intradiscal injections to the lumbar spine of rats were performed under sterile conditions and general anesthesia (2% isoflurane inhalation). Once under anesthesia, IVD levels were identified using anatomical landmarks, then a 30G needle was fed through a 20G cannula to inject NT or ZNF865 targeting CRISPRi lentiviral vectors (5ul, 10^6^-10^9^ transducing units) to the L4-L5 and L5-L6 IVDs. Animals in the naïve group underwent anesthesia, however, did not receive any injections. Following injections, rats were closely monitored for complications and given access to food and water *ad libitum*.

### Von Frey Behavioral Assessment

Mechanical pain sensitivity was measured using the Chaplan up down protocol for von Frey hairs ^76–78^. Here, rats were allowed to acclimate to a wire bottom cage for 30-45 minutes or until cage exploration and self-grooming behaviors ceased. After acclimation, von Frey monofilaments designed to bend at a known specific force (0.4, 0.6, 1, 2, 4, 6, 10, and 15 g), were pressed against the plantar region of the hind paw. Withdrawal of the paw from the monofilament resulted in the next lowest filament in the series to be tested. If no withdrawal was observed, the next highest filament in the series was tested. This paradigm was repeated until five stimulations after the first change in response, up to nine total measurements. Each hind paw was tested separately beginning with the 2g monofilament with a minimum 2 minutes between each stimulation of the same paw. Testing was performed twice prior to lentivirus injections to establish a baseline, three days prior to injections, and then bi-weekly for the duration of the 8 week experiment. Data collection was performed blinded. Paw withdrawal threshold (PWT) was calculated as described in *Chaplan et al* ^76^ for each paw, animal and group at each time point. PWT was then normalized to each animals baseline measurements, then normalized to the average of the naïve, no injection group at each time point to examine the mechanical pain sensitivity over time. Groups marked with different letters are significantly different (p<0.05).

### Hargreaves Behavioral Assessment

Thermal sensitivity was measured using the Hargreaves assay to assess pain related symptoms of peripheral neuropathy in the rats^79–82^. In this assay, rats were placed in a heated glass bottom cage and allowed to acclimate for 30-45 minutes or until cage exploration and self-grooming behaviors cease. After acclimation, a focused light source was applied from below the cage to the plantar region of the hind paw, and the time it took for animals to withdraw their paws from the heat source was recorded. This process is repeated three times per animal per hindfoot with a 5-minute rest between trails. Testing was performed twice prior to lentivirus injections to establish a baseline, three days prior to injections, and then bi-weekly for the duration of the 8 week experiment. Data collection was performed blinded. This data was then used to calculate the paw withdrawal latency (PWL) for each paw, animal, and group at each time point. PWL was then normalized to each animals baseline measurement, and then normalized to the average of the naïve, no injection group, at each time point to examine the thermal pain sensitivity over time. Groups marked with different letters are significantly different (p<0.05).

### Tissue Harvesting and Sectioning

After 8 weeks, animals were sacrificed via carbon dioxide inhalation. Immediately following sacrifice, rat lumbar spines were harvested from L3-S1 and fixed in formalin (Thermo) for 48hrs. Following fixation, samples were decalcified (Immunocal, StatLab) for 72 hours, changing decalcification solution every 24 hours. Following decalcification spine segments were embedded (Tissue-Tek TEC 5 Embedding Station, Sakura Finetek) in paraffin using melted Surgipath Formula ‘R’ paraffin pellets (Leica Biosystems). Samples were then sectioned (5um) (RM2255 Rotary Microtomes, Leica Biosystems) and placed on slides.

### Histological Assessment

Following sectioning onto slides, samples were stained for hematoxylin and eosin (H&E) and alcian blue to visualize IVD morphology. Slides were then imaged under bright-field microscope (Olympus UC50) to evaluate gross morphological differences within each group.

### Disc Height Analysis

Following histological staining with alcian blue, disc height was calculated manually in ImageJ by measuring the distance between CEPs in the center of the disc for each animal. Disc height for each animal was then normalized to the naïve group at the L4-L5 or L5-L6 levels.

### Degeneration Scoring

Following histological staining with H&E and alcian blue IVD degeneration was assessed using a previously reported semi-quantitative degeneration grading scale^83^. The combined degeneration scale, scored from 0-10, is comprised of five categories. Annulus fibrosis, border between the AF and NP, NP cellularity, NP matrix, and endplate. Each category is scored on a scale from 0-2 with a 0 indicating normal IVD morphology, and a 2 representing severe degeneration^83^.

### Immunofluorescence staining

Following tissue sectioning and mounting, slides were placed in an incubator at 42°C for 45 minutes to soften wax. Samples were then deparaffinized and hydrated by washing in Xylene (Thermo Fisher) twice and 100%, 95% and 70% EtOH (Thermo Fisher) respectively before placing in DiH_2_O. Samples were then washed in TBST. Antigen retrieval was performed by placing in 99°C citrate buffer for 30 minutes for extracellular proteins or 0.1% Triton_X for 10 minutes at room temperature for intracellular proteins. Samples were then washed in TBST before blocking with background buster solution (Innovex) for 30 min. Samples were washed again with TBST before incubating with primary antibody (Proteintech, P16INK4a 28416-1-AP, VEGFA 26157-1-AP, H2AX 10856-1-AP)(Novus Biologicals, IL-6 NB600-1131) diluted in Dako diluent overnight. The next day, samples were washed with TBST, and then incubated in secondary antibody (Proteintech, CoraLite 594 SA00013-4) diluted int Dako diluent for 1 hour at room temperature. Samples were washed in TBST then covered with Vectashield Dapi mounting media before sealing. Samples were stored at 4°C until imaging (Leica SP8 Confocal).

### Statistical Analysis

RNA-seq statistical analysis was performed and described in their respective sections (Enrichr, DESeq2, α=0.05). Statistical analysis of qRT-PCR, Luminex, β-gal, p16, p21, ZNF865, Collagen, GAG, comets, disc height and degenerative scoring were performed using JMP Pro or GraphPad Prism using a one-way ANOVA with Tukey’s post hoc analysis. Statistical analysis for proliferation and behavioral testing was performed using JMP Pro with a two-way ANOVA with Tukey’s post hoc analysis. Expression level scRNA-seq data was determined to be non-normal, significant differences between ages were tested for using Kruskal-Wallis test with Dunn’s comparisons and p<0.05 considered significant in GraphPad Prism. Simple linear regression and Pearsons’s correlation were performed on percent positivity and age with p<0.05 considered significant.

---

**Table 1:**
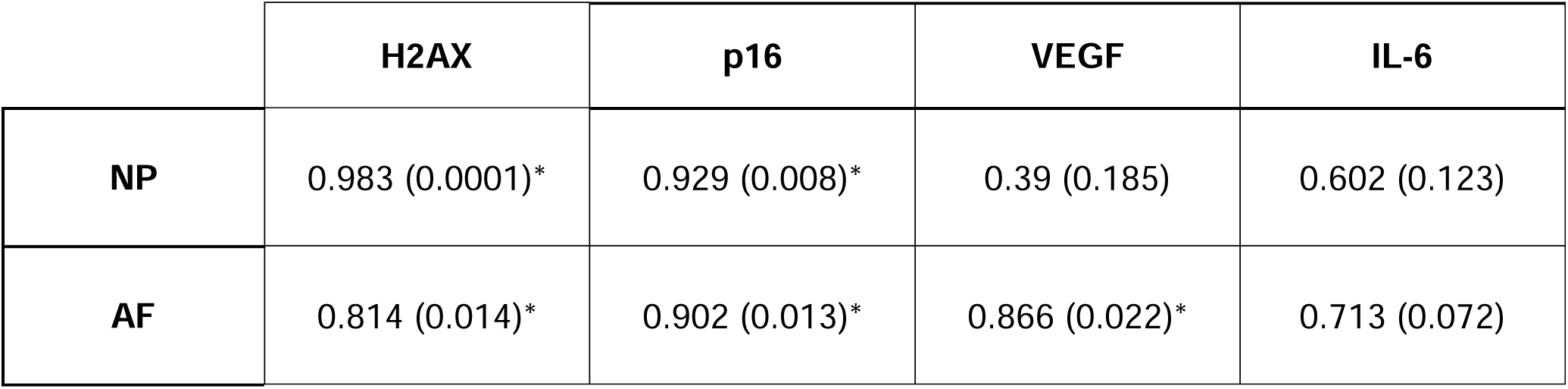
Correlation between Von Frey behavioral testing and immunofluorescence staining. Displayed as R^2^ (p-value) where *= p-value < 0.05).

|  | H2AX | p16 | VEGF | IL-6 |
| --- | --- | --- | --- | --- |
| NP | 0.983 (0.0001)* | 0.929 (0.008)* | 0.39 (0.185) | 0.602 (0.123) |
| AF | 0.814 (0.014)* | 0.902 (0.013)* | 0.866 (0.022)* | 0.713 (0.072) |

