## Supplemental materials for "ZNF865 (BLST) a Novel Regulator of DNA Damage and Cell Senescence in Back Pain"

**List of Supplementary Materials:**

Figure S1: ZNF865 is titratable in Human NP cells with CRISPRi and CRISPRa.

Figure S2: ZNF865 expression versus Thompson score in asymptomatic and symptomatic patients.

Figure S3: Secondary only staining controls for ZNF865 knockdown.

Figure S4: Correlation plots between von Frey behavioral assessment for pain and Immunofluorescence staining.

Table S1: Patient information.

Table S2: KEGG Human Genome 2021 significant pathways (From RNAseq in *HEK293* cells) ZNF865-VPR versus NT.

Table S3: KEGG Human Genome 2021 significant pathways (From RNAseq in *ASC*s). ZNF865-VPR versus NT.

Table S4: KEGG Human Genome 2021 significant pathways (From RNAseq in dhNPCs) ZNF865 VPR dhNPCs vs Non Target dhNPCs.

Table S5: ATACseq patient information.

Table S6: KEGG 2021 Human Genome significant pathways (From ATACseq) dhNPCs versus hNPCs.

Table S7: KEGG 2021 Human Genome significant pathways (From ATACseq) dhNPCs versus ZNF865-VPR dhNPCs.

Table S8: Genes associated with 70 differential chromatin accessibility sites (From ATACseq) hNPCs vs ZNF865-VPR dhNPCs.

Table S9: RNAseq patient information.

Table S10: sgRNA primers targeting the CRISPRa/i systems to ZNF865 and a nontarget control in the human for upregulation and downregulation.

Table S11:sgRNA primers targeting the CRISPRi system to ZNF865 and a nontarget control in the rat genome.

**Supplementary Materials:**

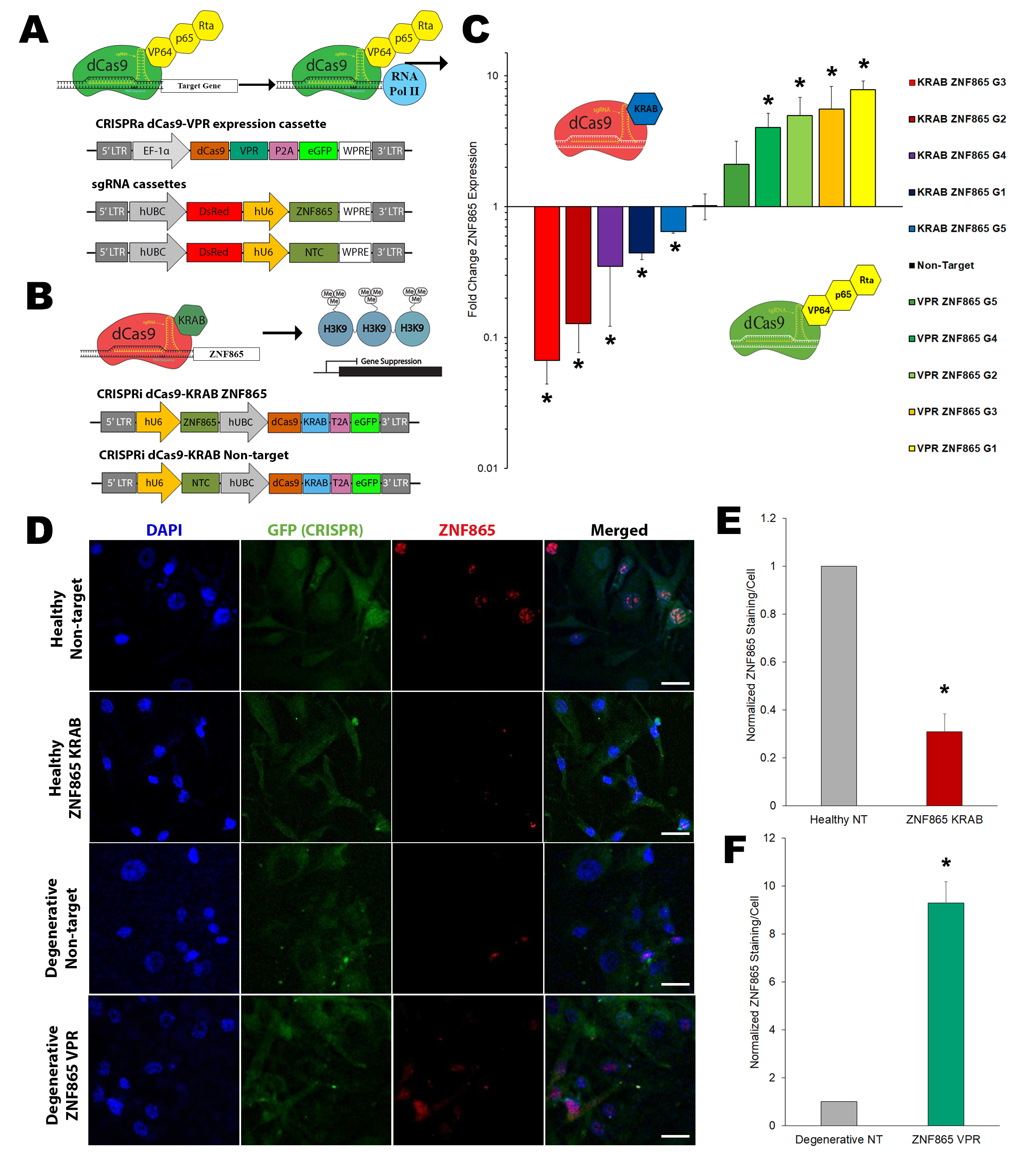

**Figure S1: ZNF865 is titratable in Human NP cells with CRISPRi and CRISPRa. A** dCas9-KRAB repression system with plasmid maps used for CRISPRi of ZNF865 or non-target. **B** dCas9-VPR activation system and plasmid maps used for CRISPRa of ZNF865 or non-target. **C** Fold change ZNF865 expression when using CRISPRi or CRISPRa systems in ASCs (n=4-5). **D** Representative immunofluorescence images after transduction with CRISPRi/CRISPRa systems with ZNF865 or non-target. **E** Normalized ZNF865 immunofluorescence staining per cell when using the CRISPRi (n=3-4) and **F** CRISPRa systems with the top performing gRNAs (n=4-5) (*Scale bars = 50 µm. significance *=p<0.05)*

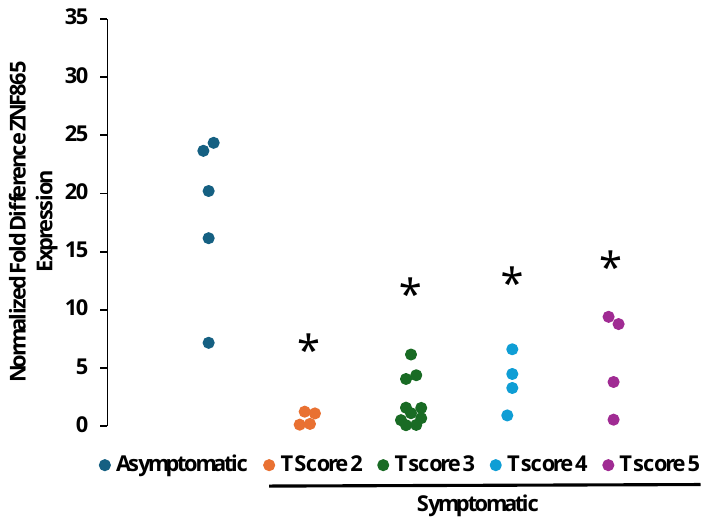

**Figure S2: ZNF865 expression versus Thompson score in asymptomatic and symptomatic patients. *** = p <0.05 compared to asymptomatic group**.**

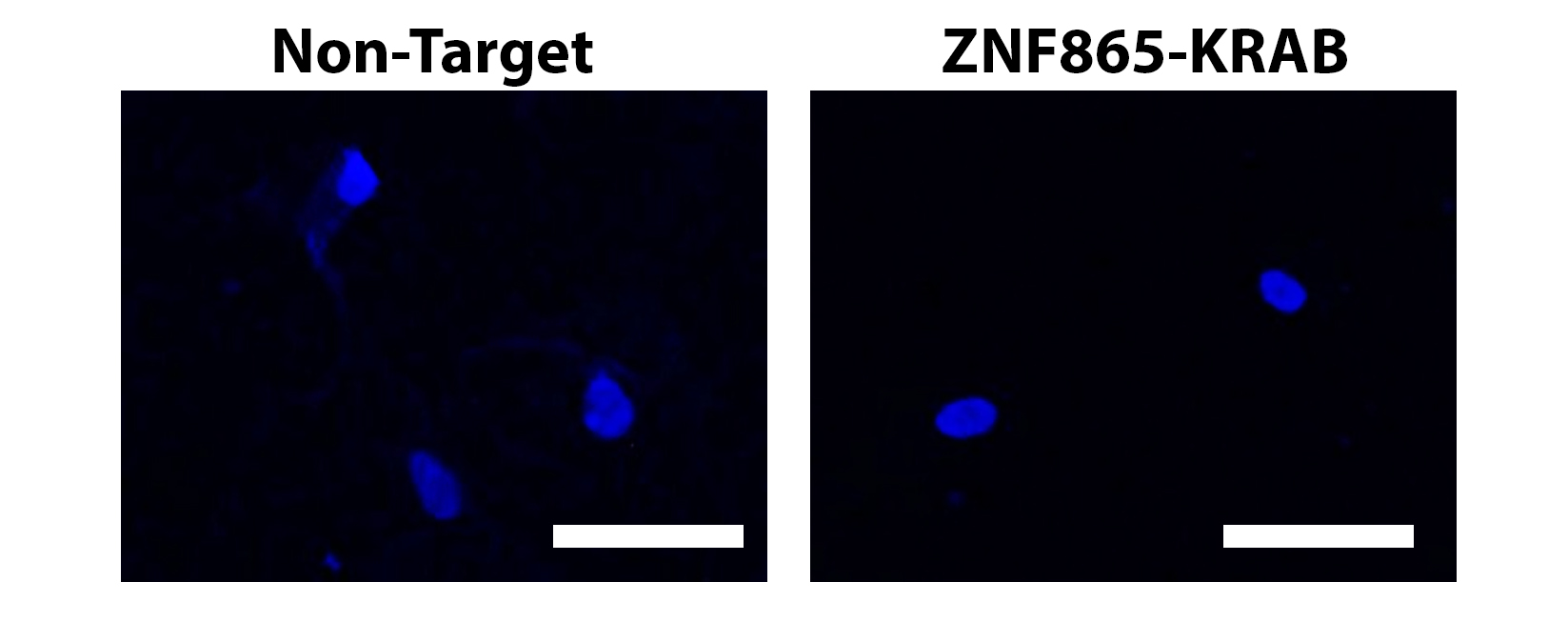

**Figure S3: Secondary only staining controls for ZNF865 knockdown.** (*Scale bars = 50 µm. significance *=p<0.05)*

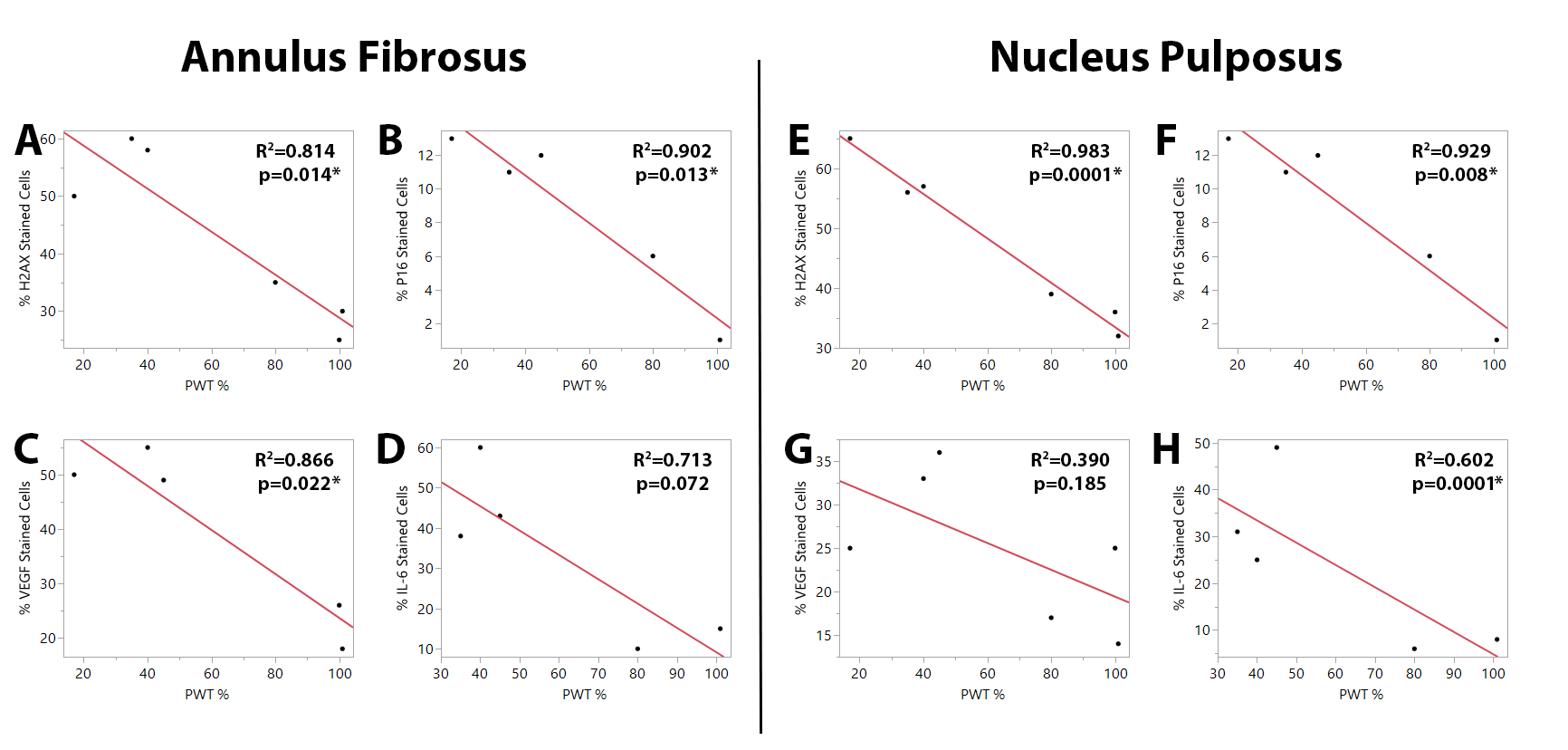

**Figure S4: Correlation plots between von Frey behavioral assessment for pain and Immunofluorescence staining.** Paw withdrawal threshold versus **A** % P16 staining, **B** % IL-6 staining, **C** %VEGF staining, **D** % H2AX staining in the nucleus pulposus. **E** % P16 staining, **F** % IL-6 staining, **G** %VEGF staining, **H** % H2AX staining in the annulus fibrosus.

**Table S1: Patient information.**

|  | T score | Gender | age |
| --- | --- | --- | --- |
| Symptomatic | 2 | M | 56 |
| Symptomatic | 2 | F | 34 |
| Symptomatic | 2 | F | 54 |
| Symptomatic | 2 | F | 54 |
| Symptomatic | 3 | M | 22 |
| Symptomatic | 3 | M | 40 |
| Symptomatic | 3 | M | 44 |
| Symptomatic | 3 | M | 57 |
| Symptomatic | 3 | M | 72 |
| Symptomatic | 3 | M | 33 |
| Symptomatic | 3 | M | 21 |
| Symptomatic | 3 | M | 59 |
| Symptomatic | 3 | F | 45 |
| Symptomatic | 3 | M | 67 |
| Symptomatic | 4 | M | 61 |
| Symptomatic | 4 | M | 39 |
| Symptomatic | 4 | M | 59 |
| Symptomatic | 4 | M | 70 |
| Symptomatic | 4 | M | 22 |
| Symptomatic | 5 | M | 50 |
| Symptomatic | 5 | F | 59 |
| Symptomatic | 5 | M | 65 |
| Symptomatic | 5 | F | 52 |
| Symptomatic | 5 | F | 54 |
| Asymptomatic | 1 | M | 61 |
| Asymptomatic | 1 | M | 31 |
| Asymptomatic | 1 | M | 38 |
| Asymptomatic | 1 | M | 28 |
| Asymptomatic | 1 | M | 35 |

**Table S2: KEGG Human Genome 2021 significant pathways (From RNAseq in *HEK293* cells) ZNF865-VPR versus NT.**

| Pathway | Adjusted p-value |
| --- | --- |
| Amyotrophic lateral sclerosis | 5.55E-20 |
| Pathways of neurodegeneration | 4.13E-18 |
| Alzheimer disease | 6.48E-17 |
| Parkinson disease | 1.02E-16 |
| Oxidative phosphorylation | 1.74E-16 |
| Huntington disease | 5.45E-16 |
| Prion disease | 3.79E-13 |
| Spliceosome | 3.55E-10 |
| Diabetic cardiomyopathy | 2.26E-09 |
| Thermogenesis | 6.03E-09 |
| Cell cycle | 1.25E-08 |
| Non-alcoholic fatty liver disease | 3.53E-08 |
| p53 signaling pathway | 5.40E-07 |
| Cellular senescence | 2.92E-06 |
| RNA transport | 4.39E-05 |
| Salmonella infection | 3.28E-04 |
| Endocytosis | 4.48E-04 |
| DNA replication | 4.99E-04 |
| Shigellosis | 7.44E-04 |
| Autophagy | 7.44E-04 |
| Ubiquitin mediated proteolysis | 0.001155331 |
| Hepatocellular carcinoma | 0.001315274 |
| Spinocerebellar ataxia | 0.001682104 |
| Bacterial invasion of epithelial cells | 0.001799459 |
| Epstein-Barr virus infection | 0.005002029 |
| Mismatch repair | 0.005002029 |
| mRNA surveillance pathway | 0.005002029 |
| mTOR signaling pathway | 0.006346221 |
| Protein processing in endoplasmic reticulum | 0.006346221 |
| Glioma | 0.007226928 |
| Chronic myeloid leukemia | 0.008454682 |
| Apoptosis | 0.010405806 |
| Basal cell carcinoma | 0.01195743 |
| RNA degradation | 0.013234801 |
| Pathogenic Escherichia coli infection | 0.013655447 |
| Viral carcinogenesis | 0.013655447 |
| Longevity regulating pathway | 0.015421792 |
| Proteoglycans in cancer | 0.015421792 |
| Colorectal cancer | 0.015421792 |
| Small cell lung cancer | 0.016643811 |
| Cardiac muscle contraction | 0.017180833 |
| Nucleotide excision repair | 0.020627196 |
| Regulation of actin cytoskeleton | 0.026414694 |
| Breast cancer | 0.026414694 |
| Propanoate metabolism | 0.026414694 |
| Various types of N-glycan biosynthesis | 0.02721576 |
| Notch signaling pathway | 0.02721576 |
| Pentose phosphate pathway | 0.029210609 |
| Hepatitis B | 0.033894794 |
| Human T-cell leukemia virus 1 infection | 0.042175878 |
| Lysine degradation | 0.048599192 |
| FoxO signaling pathway | 0.048599192 |
| Yersinia infection | 0.048599192 |
| Endometrial cancer | 0.048599192 |
| Gastric cancer | 0.048599192 |

**Table S3: KEGG Human Genome 2021 significant pathways (From RNAseq in *ASC*s). ZNF865-VPR versus NT.**

| Pathway | Adjusted P-value | Pathway | Adjusted P-value |
| --- | --- | --- | --- |
| Cell cycle | 1.76E-11 | Sphingolipid signaling pathway | 0.002706 |
| Focal adhesion | 2.90E-10 | Relaxin signaling pathway | 0.002706 |
| Pathways in cancer | 2.24E-09 | Phosphatidylinositol signaling system | 0.002836 |
| Lysosome | 2.51E-09 | Amoebiasis | 0.002874 |
| Protein processing in endoplasmic reticulum | 3.85E-09 | Progesterone-mediated oocyte maturation | 0.00323 |
| DNA replication | 3.85E-09 | AMPK signaling pathway | 0.003313 |
| Human papillomavirus infection | 3.61E-08 | Thyroid hormone signaling pathway | 0.004204 |
| Amyotrophic lateral sclerosis | 1.38E-07 | Gastric cancer | 0.004234 |
| Salmonella infection | 1.38E-07 | Oocyte meiosis | 0.004543 |
| AGE-RAGE signaling pathway in diabetic complications | 3.28E-07 | Growth hormone synthesis, secretion and action | 0.004591 |
| Cellular senescence | 3.64E-07 | Human immunodeficiency virus 1 infection | 0.004659 |
| Pathways of neurodegeneration | 5.23E-07 | Glioma | 0.005101 |
| Human T-cell leukemia virus 1 infection | 6.40E-07 | Ubiquitin mediated proteolysis | 0.005211 |
| Shigellosis | 6.90E-07 | ErbB signaling pathway | 0.005401 |
| Endocytosis | 9.00E-07 | Hypertrophic cardiomyopathy | 0.005469 |
| Adherens junction | 9.55E-07 | Gap junction | 0.006164 |
| Proteoglycans in cancer | 1.19E-06 | Choline metabolism in cancer | 0.006237 |
| Diabetic cardiomyopathy | 1.28E-06 | Renal cell carcinoma | 0.007005 |
| Regulation of actin cytoskeleton | 1.45E-06 | Vibrio cholerae infection | 0.007005 |
| TGF-beta signaling pathway | 2.08E-06 | Leukocyte transendothelial migration | 0.007454 |
| Small cell lung cancer | 2.11E-06 | Ferroptosis | 0.007454 |
| Bacterial invasion of epithelial cells | 2.11E-06 | Apelin signaling pathway | 0.007454 |
| Autophagy | 2.13E-06 | Hippo signaling pathway | 0.007543 |
| Alzheimer disease | 5.64E-06 | Ras signaling pathway | 0.007984 |
| Neurotrophin signaling pathway | 5.64E-06 | Chagas disease | 0.008369 |
| Parkinson disease | 6.83E-06 | Lysine degradation | 0.009647 |
| Axon guidance | 1.01E-05 | Lipid and atherosclerosis | 0.010094 |
| Hepatocellular carcinoma | 1.12E-05 | Breast cancer | 0.011048 |
| ECM-receptor interaction | 1.66E-05 | Phospholipase D signaling pathway | 0.013267 |
| MAPK signaling pathway | 1.91E-05 | Kaposi sarcoma-associated herpesvirus infection | 0.013474 |
| Fluid shear stress and atherosclerosis | 1.91E-05 | Epstein-Barr virus infection | 0.015663 |
| Pancreatic cancer | 2.75E-05 | Thermogenesis | 0.016597 |
| Insulin resistance | 3.30E-05 | Epithelial cell signaling in Helicobacter pylori infection | 0.016932 |
| PI3K-Akt signaling pathway | 3.64E-05 | Viral carcinogenesis | 0.017764 |
| Mitophagy | 3.64E-05 | Wnt signaling pathway | 0.018582 |
| Apoptosis | 4.55E-05 | Parathyroid hormone synthesis, secretion and action | 0.019003 |
| FoxO signaling pathway | 5.99E-05 | Melanogenesis | 0.019402 |
| Huntington disease | 6.82E-05 | Dilated cardiomyopathy | 0.019753 |
| Human cytomegalovirus infection | 1.34E-04 | Vasopressin-regulated water reabsorption | 0.020232 |
| Base excision repair | 1.53E-04 | Citrate cycle (TCA cycle) | 0.020232 |
| mTOR signaling pathway | 1.72E-04 | C-type lectin receptor signaling pathway | 0.020463 |
| Fanconi anemia pathway | 1.76E-04 | Alanine, aspartate and glutamate metabolism | 0.020463 |
| Hepatitis B | 1.96E-04 | Toxoplasmosis | 0.021236 |
| Longevity regulating pathway | 2.09E-04 | RNA degradation | 0.022609 |
| Rap1 signaling pathway | 2.18E-04 | Other types of O-glycan biosynthesis | 0.023746 |
| Spinocerebellar ataxia | 2.18E-04 | Non-alcoholic fatty liver disease | 0.024677 |
| Inositol phosphate metabolism | 2.90E-04 | Arrhythmogenic right ventricular cardiomyopathy | 0.025104 |
| Prion disease | 2.90E-04 | SNARE interactions in vesicular transport | 0.025333 |
| Estrogen signaling pathway | 3.01E-04 | Tight junction | 0.026773 |
| Pathogenic Escherichia coli infection | 3.26E-04 | Cysteine and methionine metabolism | 0.027314 |
| Homologous recombination | 4.35E-04 | Inflammatory mediator regulation of TRP channels | 0.027314 |
| RNA transport | 4.52E-04 | Regulation of lipolysis in adipocytes | 0.027371 |
| Mismatch repair | 4.52E-04 | Central carbon metabolism in cancer | 0.027371 |
| TNF signaling pathway | 4.52E-04 | RNA polymerase | 0.027371 |
| Aldosterone synthesis and secretion | 4.97E-04 | GnRH signaling pathway | 0.027371 |
| Cushing syndrome | 6.32E-04 | cGMP-PKG signaling pathway | 0.028104 |
| p53 signaling pathway | 6.32E-04 | Endometrial cancer | 0.030745 |
| Prostate cancer | 7.52E-04 | Measles | 0.03469 |
| Purine metabolism | 8.47E-04 | Pyrimidine metabolism | 0.03469 |
| Nucleotide excision repair | 9.53E-04 | Other glycan degradation | 0.034719 |
| Colorectal cancer | 9.56E-04 | Ribosome biogenesis in eukaryotes | 0.038753 |
| Phagosome | 9.56E-04 | VEGF signaling pathway | 0.038753 |
| HIF-1 signaling pathway | 0.001439 | Platelet activation | 0.039293 |
| Fc gamma R-mediated phagocytosis | 0.001482 | Signaling pathways regulating pluripotency of stem cells | 0.041146 |
| Yersinia infection | 0.001703 | mRNA surveillance pathway | 0.041518 |
| Chronic myeloid leukemia | 0.001703 | Glutathione metabolism | 0.043062 |
| Glycosaminoglycan biosynthesis | 0.001781 | Oxidative phosphorylation | 0.045129 |

**Table S4: KEGG Human Genome 2021 significant pathways (From RNAseq in dhNPCs) ZNF865 VPR dhNPCs vs Non Target dhNPCs**

| Pathway | Adjusted p-value |
| --- | --- |
| Cell cycle | 1.65E-10 |
| DNA replication | 1.68E-10 |
| Pathways in cancer | 4.12E-10 |
| Protein digestion and absorption | 1.53E-07 |
| ECM-receptor interaction | 1.04E-06 |
| Focal adhesion | 1.38E-06 |
| PI3K-Akt signaling pathway | 1.97E-05 |
| Axon guidance | 3.35E-05 |
| Basal cell carcinoma | 3.35E-05 |
| Human papillomavirus infection | 3.50E-05 |
| Breast cancer | 8.66E-05 |
| AGE-RAGE signaling pathway in diabetic complications | 9.73E-05 |
| TGF-beta signaling pathway | 1.00E-04 |
| Hippo signaling pathway | 2.31E-04 |
| Gastric cancer | 2.40E-04 |
| Wnt signaling pathway | 2.98E-04 |
| Hepatocellular carcinoma | 3.61E-04 |
| Complement and coagulation cascades | 5.53E-04 |
| Fanconi anemia pathway | 6.26E-04 |
| p53 signaling pathway | 6.26E-04 |
| Colorectal cancer | 0.001832 |
| Bladder cancer | 0.001864 |
| Proteoglycans in cancer | 0.002686 |
| Prostate cancer | 0.00292 |
| Relaxin signaling pathway | 0.003678 |
| Cushing syndrome | 0.004663 |
| Cellular senescence | 0.004987 |
| Mismatch repair | 0.005329 |
| Homologous recombination | 0.006129 |
| Human T-cell leukemia virus 1 infection | 0.007131 |
| Small cell lung cancer | 0.008761 |
| Calcium signaling pathway | 0.009037 |
| Gap junction | 0.013074 |
| Acute myeloid leukemia | 0.014962 |
| Glycine, serine and threonine metabolism | 0.016079 |
| Progesterone-mediated oocyte maturation | 0.019913 |
| Arrhythmogenic right ventricular cardiomyopathy | 0.019913 |
| Melanogenesis | 0.021449 |
| Signaling pathways regulating pluripotency of stem cells | 0.02368 |
| MAPK signaling pathway | 0.02454 |
| Malaria | 0.02454 |
| Melanoma | 0.02454 |
| Ras signaling pathway | 0.025465 |
| Oocyte meiosis | 0.026862 |
| Fluid shear stress and atherosclerosis | 0.03041 |
| Amoebiasis | 0.046035 |

**Table S5: ATACseq patient information**

| **library number** | **genotype** | **sample name** | **technical replicate** | **sex** | **age** | **sample designation** |
| --- | --- | --- | --- | --- | --- | --- |
| 1 | hNPCs | atac_hnpc_r1 | 1 | female | 69 | 23835X1 |
| 2 | dhNPCs | atac_dhnpc_r1 | 1 | male | 60 | 23835X2 |
| 3 | ZNF865-dhNPCs | atac_znf865_dhnpc_r1 | 1 | male | 32 | 23835X3 |
| 4 | hNPCs | atac_hnpc_r2 | 2 | female | 69 | 23835X4 |
| 5 | dhNPCs | atac_dhnpc_r2 | 2 | male | 60 | 23835X5 |
| 6 | ZNF865-dhNPCs | atac_znf865_dhnpc_r2 | 2 | male | 32 | 23835X6 |

**Table S6: KEGG 2021 Human Genome significant pathways (From ATACseq) dhNPCs versus hNPCs.**

| **Term** | **P-value** |
| --- | --- |
| PI3K-Akt signaling pathway | 1.02E-06 |
| Calcium signaling pathway | 2.41E-06 |
| Pathways in cancer | 5.12E-06 |
| Parathyroid hormone synthesis, secretion and action | 2.41E-05 |
| Non-small cell lung cancer | 2.46E-05 |
| Focal adhesion | 3.75E-05 |
| Vascular smooth muscle contraction | 5.02E-05 |
| Human papillomavirus infection | 5.14E-05 |
| cGMP-PKG signaling pathway | 7.27E-05 |
| Glycosaminoglycan biosynthesis | 1.02E-04 |
| Long-term potentiation | 1.24E-04 |
| MAPK signaling pathway | 1.74E-04 |
| Inflammatory mediator regulation of TRP channels | 1.95E-04 |
| Thyroid hormone signaling pathway | 2.06E-04 |
| Hippo signaling pathway | 2.72E-04 |
| Proteoglycans in cancer | 2.86E-04 |
| Insulin secretion | 2.88E-04 |
| Melanogenesis | 2.97E-04 |
| Dopaminergic synapse | 3.05E-04 |
| Longevity regulating pathway | 3.41E-04 |
| Gastric acid secretion | 5.74E-04 |
| Amphetamine addiction | 5.90E-04 |
| Ras signaling pathway | 6.60E-04 |
| Hedgehog signaling pathway | 6.98E-04 |
| GnRH signaling pathway | 7.76E-04 |
| Wnt signaling pathway | 8.31E-04 |
| Adrenergic signaling in cardiomyocytes | 8.88E-04 |
| Rap1 signaling pathway | 9.25E-04 |
| Fc gamma R-mediated phagocytosis | 0.001292806 |
| Phosphatidylinositol signaling system | 0.001292806 |
| Aldosterone synthesis and secretion | 0.001459984 |
| cAMP signaling pathway | 0.001487289 |
| Renin secretion | 0.001801153 |
| Salivary secretion | 0.002058985 |
| Basal cell carcinoma | 0.002209771 |
| Spinocerebellar ataxia | 0.002256952 |
| AMPK signaling pathway | 0.002770523 |
| Gastric cancer | 0.003845605 |
| Thyroid hormone synthesis | 0.004042767 |
| Endocrine and other factor-regulated calcium reabsorption | 0.004487966 |
| Human cytomegalovirus infection | 0.005332161 |
| Oxytocin signaling pathway | 0.005802164 |
| Cushing syndrome | 0.006278458 |
| Cellular senescence | 0.006786491 |
| Fluid shear stress and atherosclerosis | 0.007142556 |

**Table S8: Genes associated with 70 differential chromatin accessibility sites (From ATACseq) hNPCs vs ZNF865-VPR dhNPCs.**

| OR4F16 | CIRBP | GPC6 | MTRNR2L1 | RNF181 | TM4SF4 |
| --- | --- | --- | --- | --- | --- |
| OR4F29 | CMBL | GPR68 | NCR1 | RPL26L1 | TMEM106A |
| ADAMTS9 | COPE | GRAP | NIPA1 | S1PR2 | TMEM63B |
| ADGRD1 | CORO2B | GRIFIN | NKAIN3 | SASH1 | TMEM9 |
| ADI1 | COX10 | GSTM1 | NLRP2 | SETD5 | TRIM62 |
| ADRB2 | CSGALNACT1 | GTF3C1 | NPR3 | SFSWAP | TSSK1B |
| ANKLE2 | DCP2 | H2AFY | OR4N4 | SGCD | TUBB4B |
| ANP32A | DDX49 | HECTD2 | P4HB | SH2D4A | VAMP3 |
| ARGLU1 | DMAC1 | HMHB1 | PCBD2 | SLC45A1 | VAMP5 |
| ARL4D | DNMT1 | HS3ST3A1 | PCGF5 | SNTB1 | VEGFA |
| ASPH | DRD1 | HTR4 | PDE8B | STAP2 | VRK3 |
| AUTS2 | EFNA5 | IGFN1 | PDZD8 | STEAP1 | WDR41 |
| AZIN2 | ELFN1 | IL16 | PHYHIPL | STEAP2 | WWOX |
| CAMTA1 | EMX2 | IL21R | PITX1 | STXBP5 | WWTR1 |
| CATSPER3 | EPN2 | KDM4C | PPP1R2P3 | SULF1 | YIPF5 |
| CCT5 | ERGIC1 | KIZ | PPP2R1A | SUSD4 | ZNF471 |
| CHST12 | ERRFI1 | MAD1L1 | PRICKLE2 | TARS | ZNF473 |
| FSD1 | FAM13C | MSX2 | RBFA | TLE4 | GLRX |
| GANAB | FAM155A | MTBP | RNF144A | TLNRD1 | GOLGA3 |
| TLR5 |  |  |  |  |  |

**Table S7: KEGG 2021 Human Genome significant pathways (From ATACseq) dhNPCs versus ZNF865-VPR dhNPCs**

| **Term** | **P-value** |
| --- | --- |
| Hippo signaling pathway | 1.99E-05 |
| Calcium signaling pathway | 4.80E-05 |
| TGF-beta signaling pathway | 3.65E-04 |
| Non-small cell lung cancer | 4.29E-04 |

**Table S9: RNAseq patient information**

| **Library number** | **Genotype** | **Sample name** | **Biological replicate** | **Technical replicate** | **Sex** | **Age** | **Sample designation** |
| --- | --- | --- | --- | --- | --- | --- | --- |
| 1 | dhNPC | dhNPC 33 Y M | - | 1 | male | 33 | 25852X3 |
| 2 | dhNPC | dhNPC 33 Y M | - | 2 | male | 33 | 25852X4 |
| 3 | hNPC | hNPC 69 Y F | 1 | - | female | 69 | 23965X3 |
| 4 | hNPC | hNPC 32 Y F | 2 | - | female | 32 | 23965X4 |
| 5 | ZNF865-VPR dhNPC | ZNF865 VPR dhNPC 32 Y M | - | 1 | male | 32 | 23965X5 |
| 6 | ZNF865-VPR dhNPC | ZNF865 VPR dhNPC 32 Y M | - | 2 | male | 32 | 23965X6 |

**Table S10:** **sgRNA primers targeting the CRISPRa/i systems to ZNF865 and a nontarget control in the human for upregulation and downregulation.**

| **gRNAs** | **Protospacer Sequence** |
| --- | --- |
| VPR-ZNF865 Guide 1 | CCGCACAAGGATGGATGAGT |
| VPR-ZNF865 Guide 2 | GACGCCCAGAGCGTGTCGCG |
| VPR-ZNF865 Guide 3 | GAGGCGGGCATTCAAAGCGC |
| VPR-ZNF865 Guide 4 | TCGCCCACCGGAATCGGCCC |
| VPR-ZNF865 Guide 5 | ATCCTCCACGCCGGCGCCTC |
| VPR-ZNF865 Guide 6 | ACTTCCGCTTCCGGGCGGGC |
| KRAB-ZNF865 Guide 1 | ATGGGGGCGCACGACTGCTA |
| KRAB-ZNF865 Guide 2 | GACGCCCAGAGCGTGTCGCG |
| KRAB-ZNF865 Guide 3 | TCGCCCACCGGAATCGGCCC |
| KRAB-ZNF865 Guide 4 | CCATGCGGAATAGAAGCCCT |
| KRAB-ZNF865 Guide 5 | GATTATAGCGATTATATGAG |
| Non-target | TTTTTAATACAAGGTAATCT |

**Table S11:sgRNA primers targeting the CRISPRi system to ZNF865 and a nontarget control in the rat genome**

| **gRNAs** | **Protospacer Sequence** |
| --- | --- |
| Rat ZNF-865 G1 | GGGCGTGGTTCCAGTCTACG |
| Rat ZNF-865 G2 | GCGACCTGTAATAGGTCGGT |
| Rat ZNF-865 G3 | GTTCGTTCCCCCCGAAGCGC |
| Rat ZNF-865 G4 | GGCCGCTTCCGGTGGGCGAA |
| Rat ZNF-865 G5 | ACGCCGGCACAAGACACTGG |
| Rat ZNF-865 G6 | TAACGCCTGCTGCATTTAAG |
| Rat ZNF865 G7 | ACTAGGCTCCGTTCGCCCAC |
| Rat ZNF865 G8 | GGCCGCTTCCGGTGGGCGAA |
| Rat ZNF865 G9 | ATTACGCCGGCACAAGACAC |
| Rat ZNF865 G10 | CGCGACTTCCGCTTCCGGGC |
| Rat ZNF865 G11 | TATTCAGCAAGTCTGCGTTA |
| Rat ZNF865 G12 | TTAGCTGTCTATAGTGTAAT |
| Rat ZNF865 G13 | ACGTTCCTACGAATGCTCAC |
| Rat ZNF865 G14 | ATGGTATTTAACTGAAGGGT |
| Rat ZNF865 G15 | GGGGTCGAGTGGCCAGCTCG |
| Non-target | TTTTTAATACAAGGTAATCT |
